# Assessing the adaptative potential to temperature and precipitation along a steep environmental gradient in populations of European beech

**DOI:** 10.1101/2025.06.03.657567

**Authors:** Mila Tost, Ourania Grigoriadou-Zormpa, Selina Wilhelmi, Markus Müller, Henning Wildhagen, Alexandru Lucian Curtu, Oliver Gailing

**Affiliations:** Division of Plant Breeding Methodology, University of Göttingen, Carl-Sprengel-Weg 1, 37075 Göttingen (Germany); Forest Genetics and Forest Tree Breeding, University of Göttingen, Büsgenweg 2, 37077 Göttingen (Germany); Center for Integrated Breeding Research (CiBreed), University of Göttingen, Von-Siebold-Str. 4, 37075 Göttingen (Germany); Faculty of Resource Management, HAWK, Büsgenweg 1a, 37077 Göttingen (Germany); Department of Silviculture, Transilvania University of Brasov, Sirul Beethoven -1, Brașov 500123 (Romania)

**Keywords:** local adaptation, environmental association analysis, *Fagus sylvatica*, climate change

## Abstract

Climate change poses a significant threat to European beech. These concerns highlight the need to assess the adaptive potential of European beech populations to climate change. Landscape genomics, also known as environmental association analysis, is a powerful tool for identifying gene loci that contribute to local adaptation to environmental pressures. Genotypic data was collected from ∼100 adult beech trees per stand in five locations in the South-Eastern Romanian Carpathians along an altitudinal gradient associated with precipitation and temperature. In total, 53 environmental variables, comprising frost frequency change, temperature and precipitation, were extracted from the climatology data base CHELSA. Based on these variables the Ellenberg-Quotient (EQ) was calculated. We performed environmental association analysis using LFMM (latent factor mixed models) to identify Single Nucleotide Polymorphism (SNP) markers associated with environmental variables and with the principal components calculated based on these. We identified 446 SNP markers significantly associated with the first principal component (PC). These were overlapping with the SNP markers significantly associated with all environmental variables except precipitation accumulated during the growing season. The first PC was correlated with all temperature-based variables and elevation at |r| ∼0.989 to ∼0.997 and with all precipitation-based and Ellenberg-Quotient variables at |*r*| ∼0.945 to 0.950, except precipitation accumulated during the growing season. A high peak region on chromosome 2 from ∼4.56 to ∼16.27 Mb appeared in all results. This region was ∼3.47 Mb downstream from a region for local adaptation identified by Lazic et al. (2024). In this peak, 273 markers located in the coding region of 22 genes were found. Ten out these 22 were described based on a literature review. Among these ten genes, two may be involved in local adaptation based on our literature review. These two genes are *polygalacturonase QRT3-like* and *NRT1/PTR_FAMILY 5.4-like*. The gene *polygalacturonase QRT3-like* plays a role in pollen development in *Arabidopsis thaliana* L. and *Brassica rapa* L. We observed at the corresponding SNP markers, a correlation of the minor allele frequency and temperature-based environmental variables.

## Introduction

Many forest tree species are negatively affected by the direct impact of drought stress due to climate change (Hartmann et al. 2022; Piedallu et al. 2023). Published observations also report that European beech (*Fagus sylvatica* L.) shows increased mortality due to more frequent summer droughts (Martinez del Castillo et al. 2022). European beech is distributed across large parts of Europe and can be found from the south of Italy up to the south of Norway (Houston Durrant et al. 2016). It is one of the most important forest tree species in Europe, accounting for the highest percentage of broadleaf growing stock on the continent (Houston Durrant et al. 2016; FAO 2020; Rukh et al. 2023). Martinez del Castillo et al. (2022) report that the 1°C increase in temperature from 1955-1985 to 1986-2016 led to reduced beech tree growth at higher altitudes in Central Europe, such as along the Carpathians. Hartmann et al. (2022) describe that European beech forests in central Germany are suffering due to increased late frost events in montane regions. Projections assume a decline in the environmental suitability of beech in south and central Germany (Baumbach et al. 2019) and southern Europe (Martinez del Castillo et al. 2022).

Hence, the importance of studying the adaptive potential of forests to changing climatic conditions by evaluating adaptive loci and their association with environmental variables is increasing (Neale and Kremer 2011). Landscape genomics also deemed as environmental association analysis (EAA) aims to identify loci involved in local adaptation and associated with environmental variables (Frichot et al. 2013; Berg and Coop 2014; Rellstab et al. 2015). Many methods are available for EAA which comprise Bayenv (Berg and Coop 2014) and latent factor mixed models (LFMM) (Frichot et al. 2013; Frichot and François 2015). Both methods report low rates of false positive observations in comparison to other EAA methods, because they include correction methods for population structure (Frichot et al. 2013; De Villemereuil et al. 2014). Most EAA methods have difficulties to distinguish effects of local adaptation from genetic drift and demographic history (Hancock et al. 2008). Neutral markers are then used as a null distribution (Hancock et al. 2008). So-called neutral markers comprise intergenic markers which are not in close proximity to genes and are therefore assumed to only capture genetic variation due to genetic drift and demographic history, but not due to adaptation (Hancock et al. 2008). Neutral markers are, however, difficult to identify and intergenic regions do not necessarily only capture genetic variation due to genetic drift and demographic history (Frichot et al. 2013). This is even more difficult with sequencing methods like single primer enrichment technology (SPET) that are designed in particular for SNPs within and close to genes (Scaglione et al. 2019). LFMM addresses this problem of confounding effects of genetic drift and demographic history by testing the correlation between environmental and genetic variation while estimating the effects of hidden factors such as residual levels of population structure (Frichot et al. 2013). EAA methods return SNP markers which are associated with tested environmental variables (Frichot et al. 2013; Rellstab et al. 2015). Identified SNPs could help to detect genes or loci involved in adaptation to specific environmental pressures and be compared to other studies. Previous studies in European beech investigated adaptation to different environments with *F_ST_* outlier approaches (Csilléry et al. 2014; Krajmerová et al. 2017; Cuervo-Alarcon et al. 2018). Some more recent studies used LFMM and other EAA methods but did not work with the chromosome-level genome assembly of European beech (Postolache et al. 2021; Müller et al. 2024). The chromosome-level genome assembly of European beech enabled us to perform a comprehensive analysis of the identified genes. Only few studies used LFMM and other EAA methods and worked with SNP marker data sets based on the chromosome-level genome assembly of European beech (Lazic et al. 2024).

Sampling for EAA studies is often done along environmental gradients (Rellstab et al. 2015). For this study, we used 497 individual trees from five European beech stands in the South-Eastern Romanian Carpathians sampled along an altitudinal gradient ranging from 550 up to 1400 m above sea level associated with precipitation and temperature. In total, 53 environmental variables were investigated separately in EAA by using LFMM. Additionally, we also performed a principal component analysis (PCA) on all environmental variables. Performing a PCA on environmental variables can help to reduce the data set and to decrease the number of tests (Rellstab et al. 2015). Though, the interpretation of the principal components (PCs) can be difficult and true positives may be missed, leading to misclassifications (Rellstab et al. 2015). It is recommended to only run EAA on environmental PCs when they can be interpreted (Rellstab et al. 2015).

The climatic conditions in the summer of the South-Eastern Romanian Carpathians may be similar to the future climatic conditions in mountainous regions of Germany (Baumbach et al. 2019; Martinez del Castillo et al. 2022). Patterns of fine-scale local adaptation to different environmental pressures will be described along a steep environmental gradient in these populations of European beech. Furthermore, environmental variables involved in local adaptation will be identified. Results from this study can help in the adaptive management of European and German beech forests, but there is still a remaining uncertainty regarding the future climatic conditions (Jandl et al. 2019).

## Material and methods

### Data sampling and stand descriptions

The used data set consists of 497 individual trees collected from five beech stands in the South-Eastern Romanian Carpathians along an altitudinal gradient associated with precipitation and temperature. The beech stands are referred to as Lempes, Tampa, Solomon, Lupului and Ruia. In each stand, approximately 100 trees were sampled in August 2021 for DNA analyses. Lempes was the lowest elevation site at 550 to 600 m, followed by Tampa located at 650-700 m, Solomon at 800 to 900 m and Lupului at 1000 to 1150 m. Ruia was the highest location at 1300 to 1450 m. The stands are naturally regenerating and were established between 100 to 160 years ago. Ruia and Lupului were established 150 and 160 years ago. Solomon, Tampa and Lempes were established 110 to 130, 130 to 150 and 100 years ago (Grigoriadou-Zormpa et al. 2024). The diameter at breast height (DBH) in Ruia is on average 35.69 cm, in Lupului 52.52 cm, in Solomon 29.52 cm, in Tampa 41.28 cm and in Lempes 47.51 cm. The distribution of DBH measurements across the different stands is available in supplemental Figure S1. For further details see Grigoriadou-Zormpa et al. (2024), in which also fine-scale spatial genetic structure of the stands is reported.

### Genotypic data

Individual trees were sequenced by using the single primer enrichment technology (SPET) (Scaglione et al. 2019). This sequencing method targets SNPs located within and close to genes and flanking regions of ±2 kb (Scaglione et al. 2019). The target regions were determined by previous whole-genome sequencing of a subset of 96 individual trees by IGATech (IGA Technology Services Srl, Udine, Italy). Reads were aligned to the *Fagus sylvatica* L. reference genome version 2 (Mishra et al. 2022) by using BWA-MEM v0.7.17 (Li and Durbin 2009). Only aligned reads with a mapping quality ≥10 were kept and duplicated reads were removed. Variant calling was done along with filtering for minimum read number of an allele in a sample with Freebayes v1.3.6 (Garrison and Marth 2012; Grigoriadou-Zormpa et al. 2024). Normalization and filtering for raw read depth were performed using bcftools (Danecek et al. 2011; Grigoriadou-Zormpa et al. 2024). After this, a total of 838,522 SNP markers were kept, which were used in the downstream analysis.

### Environmental data

In total, 53 environmental variables were used in this analysis. The environmental variables comprise recorded weather data from 1981 until 2010. Environmental data was downloaded from the climatology data base CHELSA v2.1 (Karger et al. 2017; Karger et al. 2018; Karger et al. 2020a; Karger et al. 2020b; Beck et al. 2020; Brun et al. 2022). The environmental variables comprised elevation in m above sea level, frost change frequency (fcf), and precipitation and temperature variables (Karger et al. 2017; Karger et al. 2018; Karger et al. 2020a; Karger et al. 2020b; Beck et al. 2020; Brun et al. 2022). All these variables were available for groups of individual trees, because the coordinates of the trees were recorded and the climatology data base CHELSA v2.1 has a very high resolution (30 arc sec, ∼1km).

Additionally, the Ellenberg-Quotient was calculated as 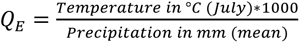 (Ellenberg 2009). Precipitation was modelled as monthly precipitation from January to December, mean annual precipitation and the precipitation accumulated over the growing season (gsp) (Karger et al. 2017; Karger et al. 2018; Karger et al. 2020a; Karger et al. 2020b; Beck et al. 2020; Brun et al. 2022). The temperature was measured as mean, minimum and maximum daily air temperature from January to December (Karger et al. 2017; Karger et al. 2018; Karger et al. 2020a; Karger et al. 2020b; Beck et al. 2020; Brun et al. 2022). The distribution of environmental variables across stands is shown in supplemental Figure S2.

We examined the correlations of the environmental variables with each other, which are shown in supplemental Figure S3. Correlations were analyzed using R version 4.3.1 (R Core Team 2024). A principle component analysis (PCA) was performed on all 53 environmental variables by using the *prcomp()* R function in the R version 4.3.1 (R Core Team 2024). The first 14 principal components (PCs) with the largest eigenvalues were also investigated using LFMM. Supplemental Figure S4 shows the two first PCs of the PCA on the environmental variables and the eigenvalues of all PCs.

### Environmental association analysis (EAA)

To test for associations between SNPs and environmental gradients, we used the latent factor mixed models (LFMM) (Frichot et al. 2013) implemented in LEA (Landscape and Ecological Association Studies) R package (R Core Team 2024; Frichot and François 2015). This LFMM model is an improved version of the LFMM algorithm from Frichot et al. (2013), which implements LFMM based on a Bayesian bootstrap approach (Frichot and François 2015). LFMM detects correlations between environmental variables and genetic variation while considering random effects due to population structure (Frichot et al. 2013). LFMM estimates confounders also referred to as latent factors (Frichot et al. 2013; Frichot and François 2015). Latent factors are then included in a statistical model for testing associations between genotypes and the environmental variable (Frichot et al. 2013; Frichot and François 2015). To run LFMM, missing data had to be imputed (Frichot et al. 2013; Frichot and François 2015), after removal of markers with a missingness >0.05 (see supplemental Fig. S5). Imputation was then done with the *impute()* function from the LEA R package (R Core Team 2024; Frichot and François 2015). LFMM tested all 53 environmental variables and 14 environmental PCs separately (Frichot et al. 2013; Frichot and François 2015). The number of latent factors was set to three based on the results of the population structure analysis (see Fig. 1 and below). LFMM results were then Benjamini-Hochberg (BH) corrected (Benjami and Hochberg 1995). BH correction was conducted with FDRestimation R package (Murray and Blume 2021). To determine whether the significance thresholds are appropriate, the number of false-positive observations exceeding these thresholds was determined by permutation tests. For these permutation tests, the first environmental PC was used. The permutation was performed while maintaining the population or stand structure, with the order of the stands randomized. The stands were randomized by changing the order along the altitudinal gradient to remove association with the altitudinal gradient. This randomization scheme is shown in supplemental Table S6. Additionally, the order of individuals within the stands was randomized by using the *sample()* R function in the R version 4.4.0 (R Core Team 2025). A total of ten replication schemes with ten replications each were carried out. The results of the randomizations are shown in supplemental Table S6. For the significance thresholds based on the BH correction, an average of ∼39 false-positive results were observed for the ten different permutations with ten replications (supplementary Table S6).

**Fig. 1:**
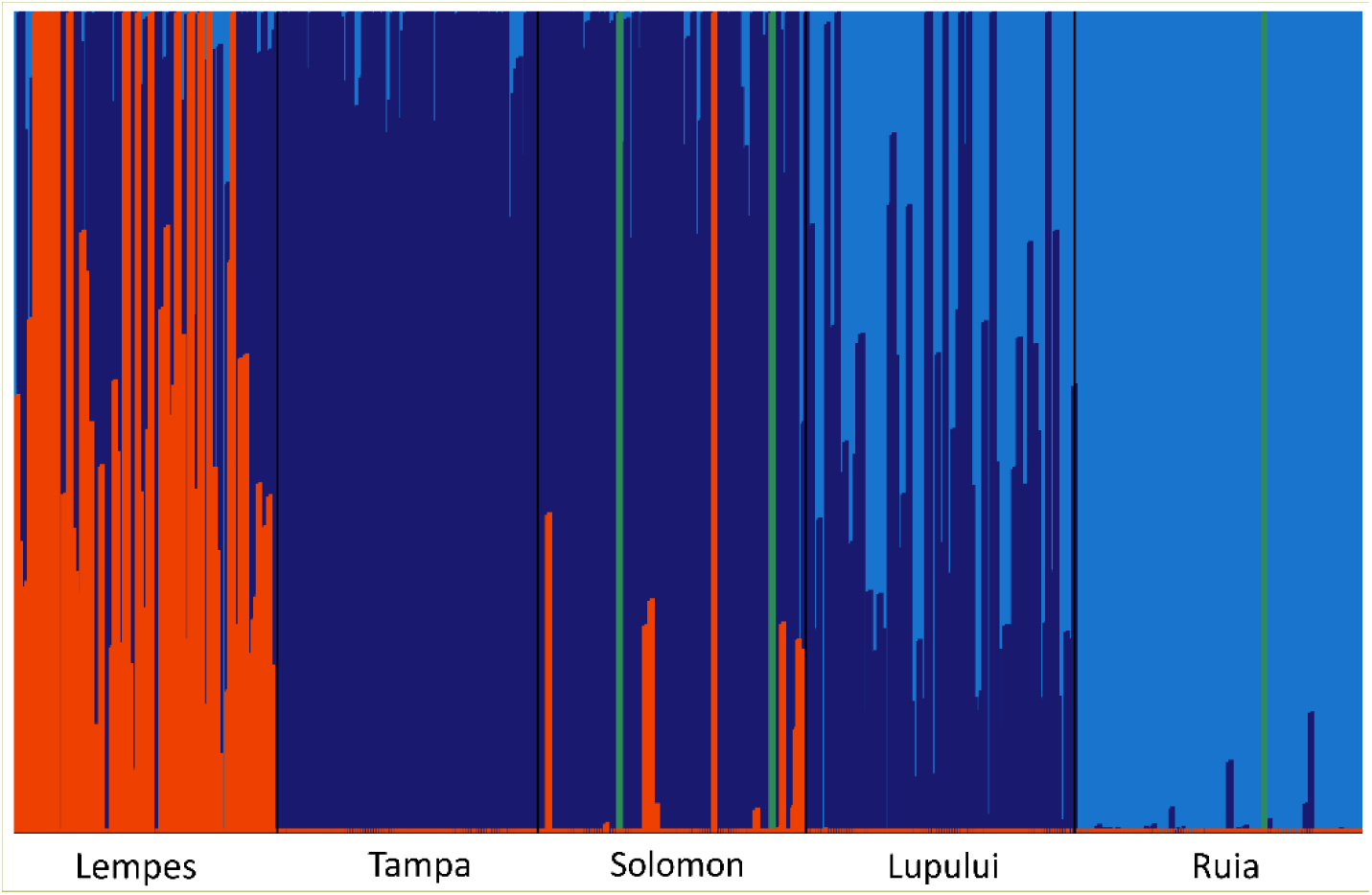
Population structure based on fastStructure (Raj et al. 2014) of the analyzed five beech stands Lempes, Tampa, Solomon, Lupului and Ruia. The different colors represent the identified clusters based on the population structure analysis.

### Assessing population stratification and assesment of confounders

To interfer population structure and to identify the number of confounders in the data set, the fastStructure algorithm was used (Raj et al. 2014). FastStructure uses variational Bayesian inference methods to determine the underlying ancestry proportions, similar to Structure (Pritchard et al. 2000), but faster and with a more flexible priority distribution (Raj et al. 2014). FastStructure achieves similar accuracies comparable to ADMIXTURE (Raj et al. 2014). FastStructure was run in python version 3.11.6 (Van Rossum and Drake 2009). The results were analyzed using R version 4.3.1 (R Core Team 2024). Results of the population structure analysis are shown in Figure 1. There is a clear, albeit slight, differentiation between the stands showing different proportions of three main genetic clusters (Fig. 1).

Additionally, we conducted an analysis of molecular variance (AMOVA) to assess the variation among stands and within stands with the poppr R package (Kamvar et al. 2013). According to the analysis of molecular variance (AMOVA) the variation among stands was 3.11% and between samples within stands was 96.89% (see supplemental Table S7). The pairwise *F_ST_* matrix is shown in supplemental Table S8. The pairwise *F_ST_* matrix was calculated based on the StAMPP R package (Pembleton et al. 2013).

## Results

The first environmental principal component (PC) explained 9.5% of the total variance and the first two PCs accounted together for 15% (see supplemental Fig. S3). Nonetheless, the LFMM results based on these PCs look very similar, and we observed that additional LFMM analysis on subsequent environmental PCs did not yield additional significant associations (see supplemental Fig. S9). The first environmental PC was correlated at |*r*| ∼0.989 to ∼0.997 with all temperature-based variables and elevation and at |*r*| ∼0.950 to ∼0.945 with all precipitation-based variables and EQ. The correlation coefficients with frost frequency change (fcf) and precipitation accumulated over the growing season (gsp) were |*r*| ∼0.813 and ∼0.692.

Figure 2 shows the results of the LFMM analysis of the first environmental PC with Benjamini-Hochberg (BH) corrected p-values on the –log_10_ scale. Environmental PC1 was significantly associated with 446 markers, of which 288 are located on chromosome 2 (Fig. 2). A high peak region appears on chromosome 2 from ∼4.56 to ∼16.27 Mb, in which 273 markers are located. The LFMM analysis of the individual environmental variables shows a similar pattern except for gsp (see supplemental Fig. S10).

**Fig. 2:**
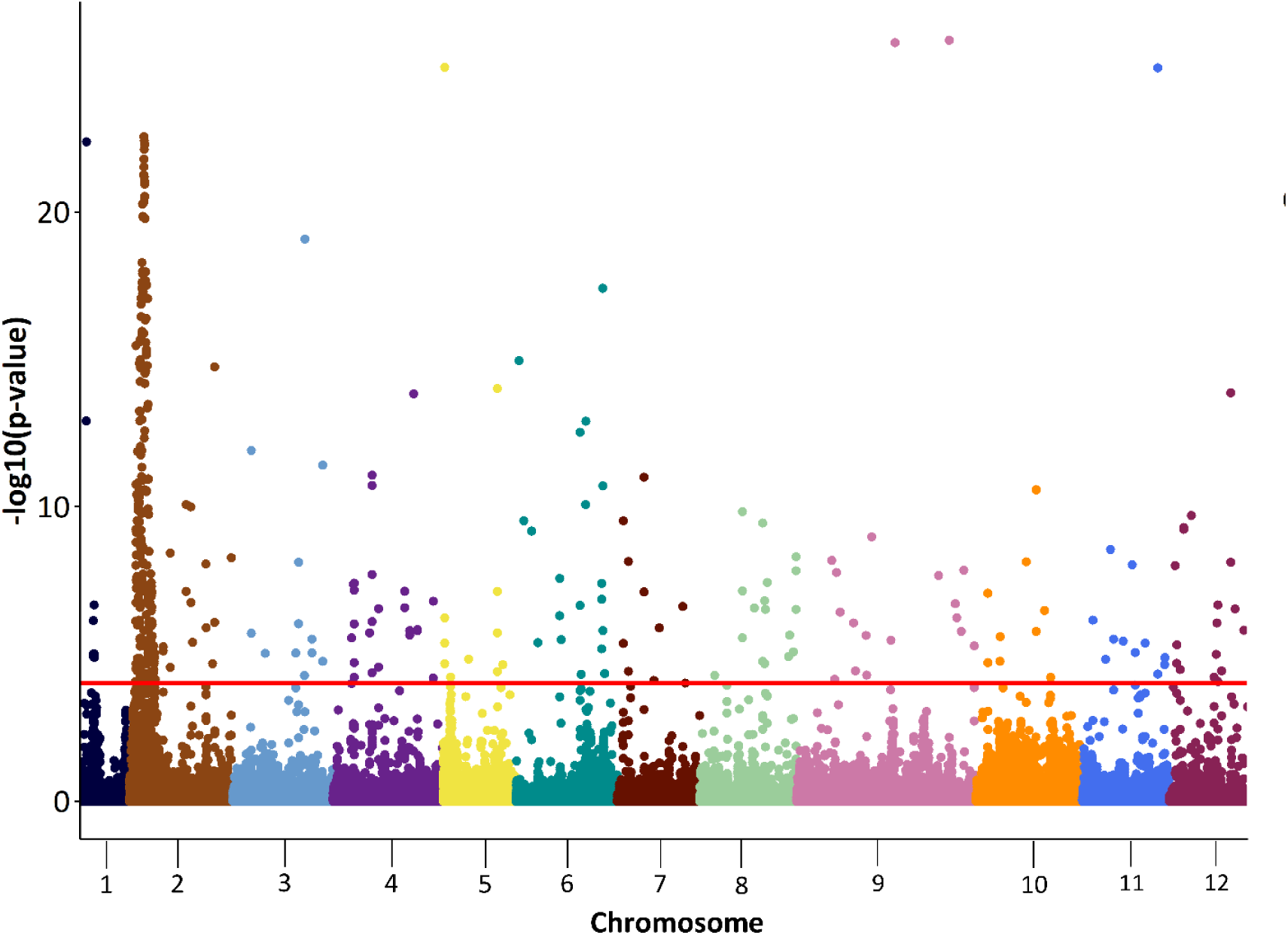
Manhattan plots based on LFMM results for environmental PC1 with Benjamini-Hochberg (BH) corrected p-values on the –log_10_ scale. The red horizontal line indicates significance thresholds with p ≤0.0001.

Figure 3A shows the LFMM results with Benjamini-Hochberg (BH) corrected p-values on the –log_10_ scale only for chromosome 2. The pairwise linkage disequilibrium (LD) heatmap was calculated for the high peak region on chromosome 2 from ∼4.56 to ∼16.27 Mb (Fig. 3C). The blue vertical line shows the region for local adaptation observed in a study by Lazic et al. (2024) from 0.79 Mb to 1.09 Mb on chromosome 2. The coding regions of all gene variants observed across this region are shown in Figure 3B. The pairwise LD was 0.2586 in Ruia, 0.2503 in Lupului, 0.2687 in Solomon, 0.2620 in Tampa and 0.2668 in Lempes (Fig. 3C).

**Fig. 3:**
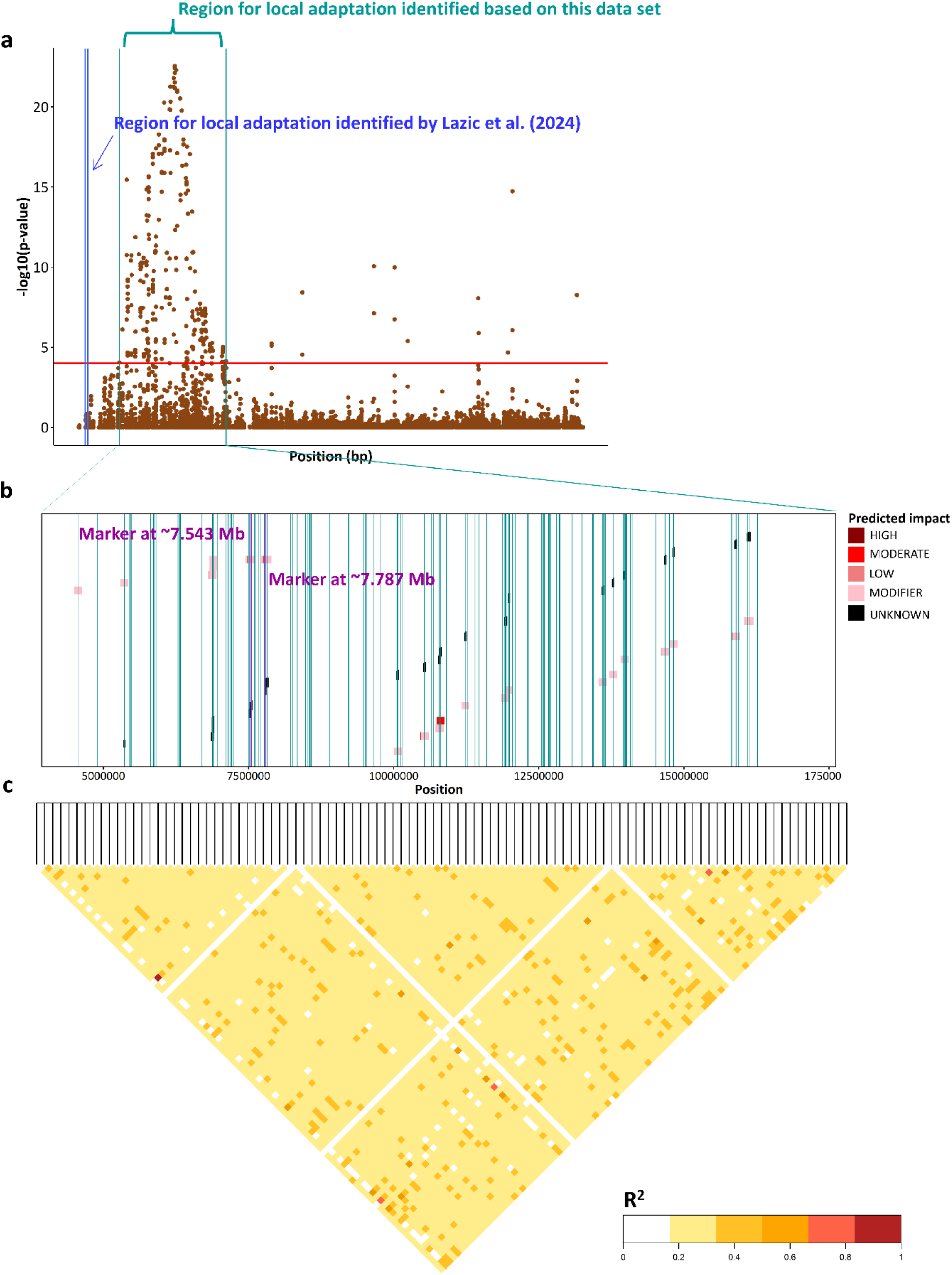
High peak region on chromosome 2 from ∼4.56 to ∼16.27 Mb with **a**) Benjamini-Hochberg (BH) corrected p-values on the –log_10_ scale (red horizontal line indicates significance thresholds with p ≤0.0001), the blue vertical line shows the region for local adaptation observed in Lazic et al. (2024), **b**) the coding regions of underlying genes and significant markers (green vertical lines) and **c**) the LD heat map in this region for Solomon.

Table 1 shows all significant SNP associations on chromosome 2 within the high peak region. In total, we observed 237 SNP markers located between ∼4.56 Mb and ∼16.27 Mb. Some markers were located within coding regions of 22 different genes, many markers were located within the coding region of the same gene, and some markers were outside of the coding regions of genes (Fig. 3). In total, we found ten described genes based on our literature review (Table 1).

**Table 1:**
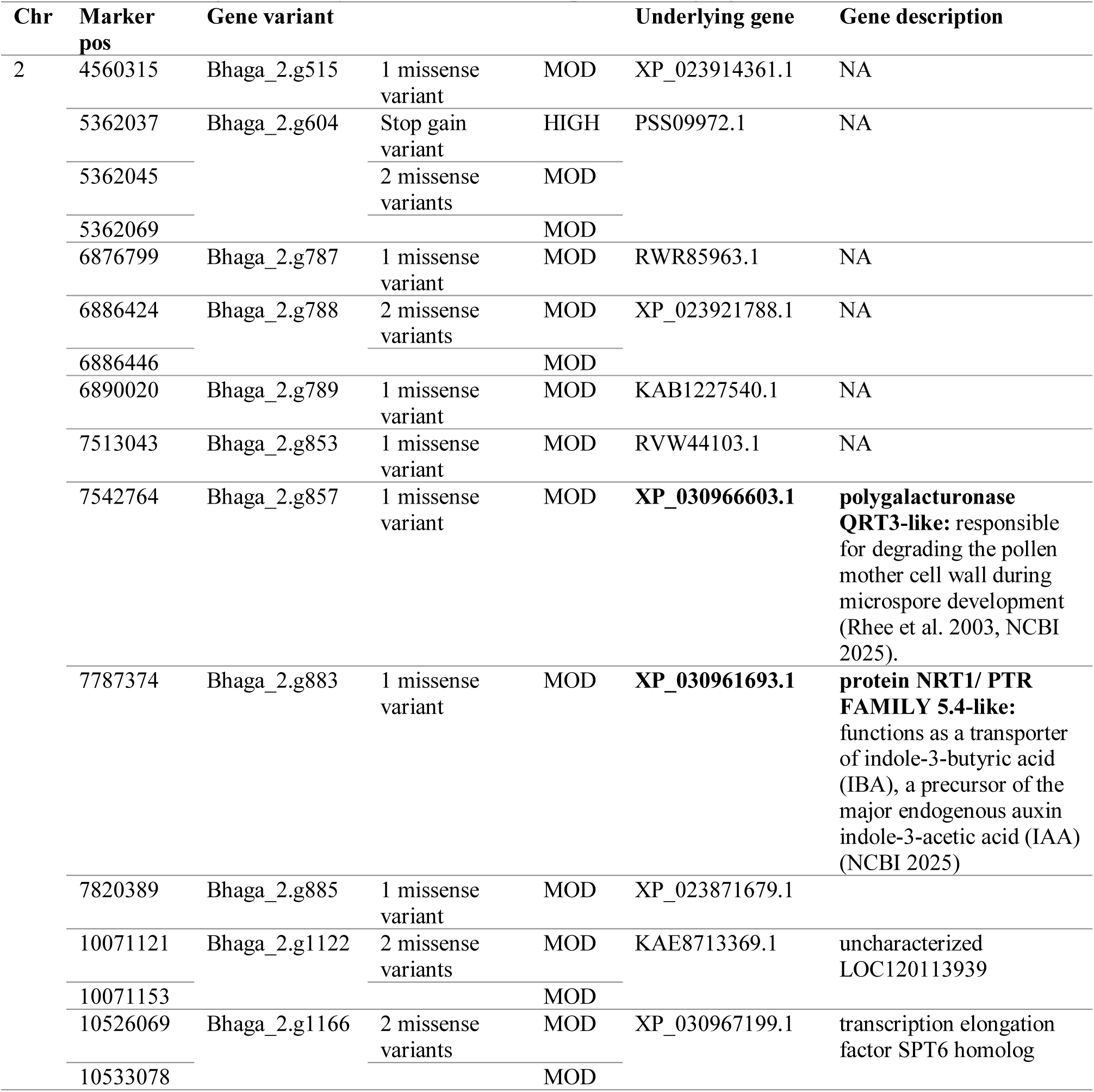
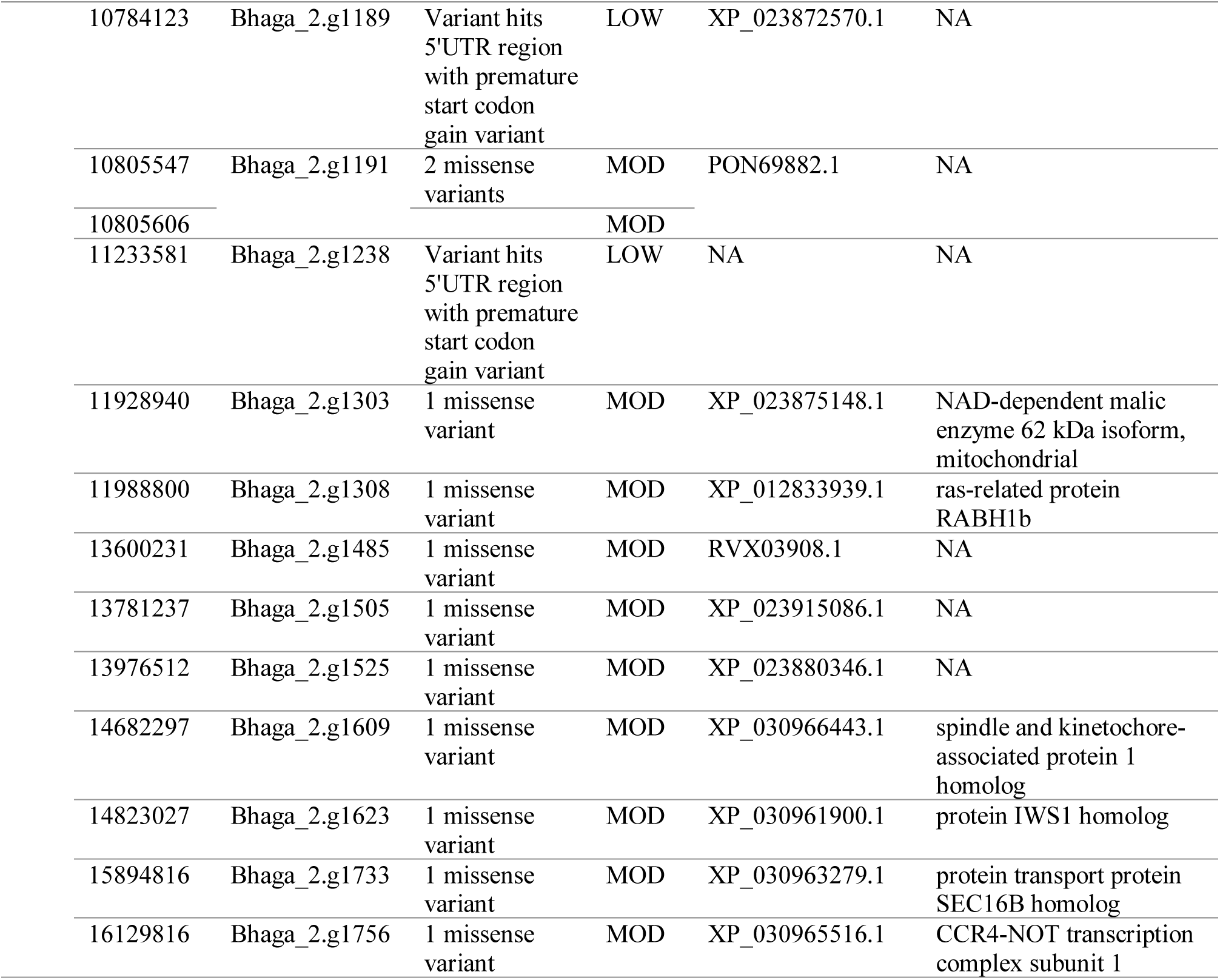
List of candidate SNPs with gene annotations associated with environmental PC1 with Bonferroni-Hochberg corrected p-values at a significance levels of p ≤0.0001 and their annotated gene variants, the underlying genes with their locations, functions and predicted impact with HIGH, MOD (moderate) and LOW based on SnpEff (Cingolani et al. 2012). Candidate genes that may be involved in local adaptation are highlighted in bold face.

| Chr | Marker pos | Gene variant |  |  | Underlying gene | Gene description |
| --- | --- | --- | --- | --- | --- | --- |
| 2 | 4560315 | Bhaga_2.g515 | 1 missense variant | MOD | XP_023914361.1 | NA |
|  | 5362037 | Bhaga_2.g604 | Stop gain variant | HIGH | PSS09972.1 | NA |
|  | 5362045 |  | 2 missense variants | MOD |  |  |
|  | 5362069 |  |  | MOD |  |  |
|  | 6876799 | Bhaga_2.g787 | 1 missense variant | MOD | RWR85963.1 | NA |
|  | 6886424 | Bhaga_2.g788 | 2 missense variants | MOD | XP_023921788.1 | NA |
|  | 6886446 |  |  | MOD |  |  |
|  | 6890020 | Bhaga_2.g789 | 1 missense variant | MOD | KAB1227540.1 | NA |
|  | 7513043 | Bhaga_2.g853 | 1 missense variant | MOD | RVW44103.1 | NA |
|  | 7542764 | Bhaga_2.g857 | 1 missense variant | MOD | **XP_030966603.1** | **polygalacturonase QRT3-like:** responsible for degrading the pollen mother cell wall during microspore development (Rhee et al. 2003, NCBI 2025). |
|  | 7787374 | Bhaga_2.g883 | 1 missense variant | MOD | **XP_030961693.1** | **protein NRT1/ PTR FAMILY 5.4-like:** functions as a transporter of indole-3-butyric acid (IBA), a precursor of the major endogenous auxin indole-3-acetic acid (IAA) (NCBI 2025) |
|  | 7820389 | Bhaga_2.g885 | 1 missense variant | MOD | XP_023871679.1 |  |
|  | 10071121 | Bhaga_2.g1122 | 2 missense variants | MOD | KAE8713369.1 | uncharacterized LOC120113939 |
|  | 10071153 |  |  | MOD |  |  |
|  | 10526069 | Bhaga_2.g1166 | 2 missense variants | MOD | XP_030967199.1 | transcription elongation factor SPT6 homolog |
|  | 10533078 |  |  | MOD |  |  |

|  |  |  |  |  |  |
| --- | --- | --- | --- | --- | --- |
| 10784123 | Bhaga_2.g1189 | Variant hits<br>5'UTR region<br>with premature<br>start codon<br>gain variant | LOW | XP_023872570.1 | NA |
| 10805547 | Bhaga_2.g1191 | 2 missense<br>variants | MOD | PON69882.1 | NA |
| 10805606 |  |  | MOD |  |  |
| 11233581 | Bhaga_2.g1238 | Variant hits<br>5'UTR region<br>with premature<br>start codon<br>gain variant | LOW | NA | NA |
| 11928940 | Bhaga_2.g1303 | 1 missense<br>variant | MOD | XP_023875148.1 | NAD-dependent malic<br>enzyme 62 kDa isoform,<br>mitochondrial |
| 11988800 | Bhaga_2.g1308 | 1 missense<br>variant | MOD | XP_012833939.1 | ras-related protein<br>RABH1b |
| 13600231 | Bhaga_2.g1485 | 1 missense<br>variant | MOD | RVX03908.1 | NA |
| 13781237 | Bhaga_2.g1505 | 1 missense<br>variant | MOD | XP_023915086.1 | NA |
| 13976512 | Bhaga_2.g1525 | 1 missense<br>variant | MOD | XP_023880346.1 | NA |
| 14682297 | Bhaga_2.g1609 | 1 missense<br>variant | MOD | XP_030966443.1 | spindle and kinetochore-<br>associated protein 1<br>homolog |
| 14823027 | Bhaga_2.g1623 | 1 missense<br>variant | MOD | XP_030961900.1 | protein IWS1 homolog |
| 15894816 | Bhaga_2.g1733 | 1 missense<br>variant | MOD | XP_030963279.1 | protein transport protein<br>SEC16B homolog |
| 16129816 | Bhaga_2.g1756 | 1 missense<br>variant | MOD | XP_030965516.1 | CCR4-NOT transcription<br>complex subunit 1 |

For the markers on chromosome 2 at ∼ 7.543 Mb and at ∼ 7.787 Mb, we found two interesting underlying candidate genes that may be involved in local adaptation. At both markers, the minor allele frequency (MAF) decreases with increasing minimum daily temperature in October following the altitudinal gradient (Fig. 4). For minimum daily temperature in October, we observed the highest absolute correlation coefficients of |*r|* ∼0.9055 (p-value = 0.03) and ∼0.9261 (p-value = 0.024). Significant correlation coefficients of the MAF and the other environmental variables are shown in supplemental Figure S11. For all temperature-based variables, frost change frequency (fcf) and Ellenberg- Quotient (EQ), we observed the same trend (see supplemental Fig. S11 and Table S12). For all precipitation-based variables and elevation, we observed the opposite trend (see supplemental Fig. S11 and Table S12). The correlation coefficients of the MAF and precipitation-based variables, EQ and fcf were not significant. The significant correlation coefficients ranged between |*r*| ∼0.8786 and ∼0.9055 for the marker on chromosome 2 at ∼ 7.543 Mb. For the marker on chromosome 2 at ∼ 7.787 Mb, the significant correlation coefficients ranged between |*r*| ∼0.9090 and 0.9261.

**Fig. 4:**
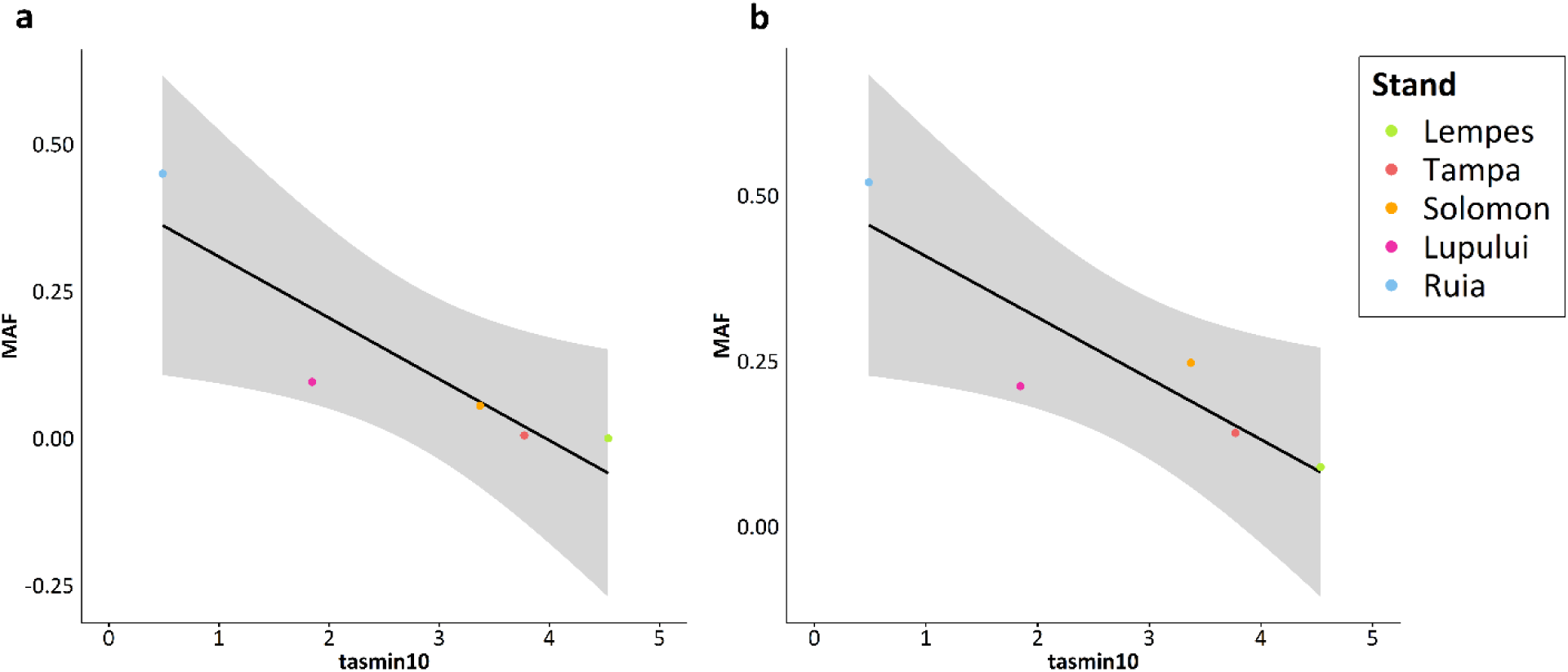
**a**) Correlation between minimum daily temperature in October (tasmin10) and the minor allele frequency (MAF) of the significant markers (p ≤0.0001) on chromosome 2 at ∼ 7.543 Mb with a correlation coefficient of -0.9055 (p-value = 0.03). This marker was annotated with gene variant *Bhaga_2.g857*, with the underlying gene *polygalacturonase QRT3-like*. **b**) Correlation between minimum daily temperature in October and the minor allele frequency (MAF) of the significant markers (p ≤0.0001) on chromosome 2 at ∼ 7.787 Mb with a correlation coefficient of -0.9261 (p-value = 0.024). This marker was annotated with gene variant *Bhaga_2.g883*, with the underlying gene protein *NRT1/ PTR FAMILY 5.4-like*.

For all other markers on chromosome 2 from ∼4.56 to ∼16.27 Mb, we observed the same trend (see supplemental Fig. S13). In all cases, the minor allele frequency (MAF) is correlated with minimum daily temperature in October (see supplemental Fig. S13). Additionally, a linear regression of the different genotypes observed at the SNP markers on chromosome 2 at ∼ 7.543 Mb (*Bhaga_2.g857*, with the underlying gene *polygalacturonase QRT3-like*) and ∼7.787 Mb (*Bhaga_2.g883*, with the underlying gene protein *NRT1/ PTR FAMILY 5.4-like*) and minimum daily temperature in October is shown in Figure 5. The regression coefficients for SNP markers on chromosome 2 at ∼ 7.543 Mb and at ∼7.787 Mb are 0.3468 and 0.1646, respectively (Fig. 5).

**Fig. 5:**
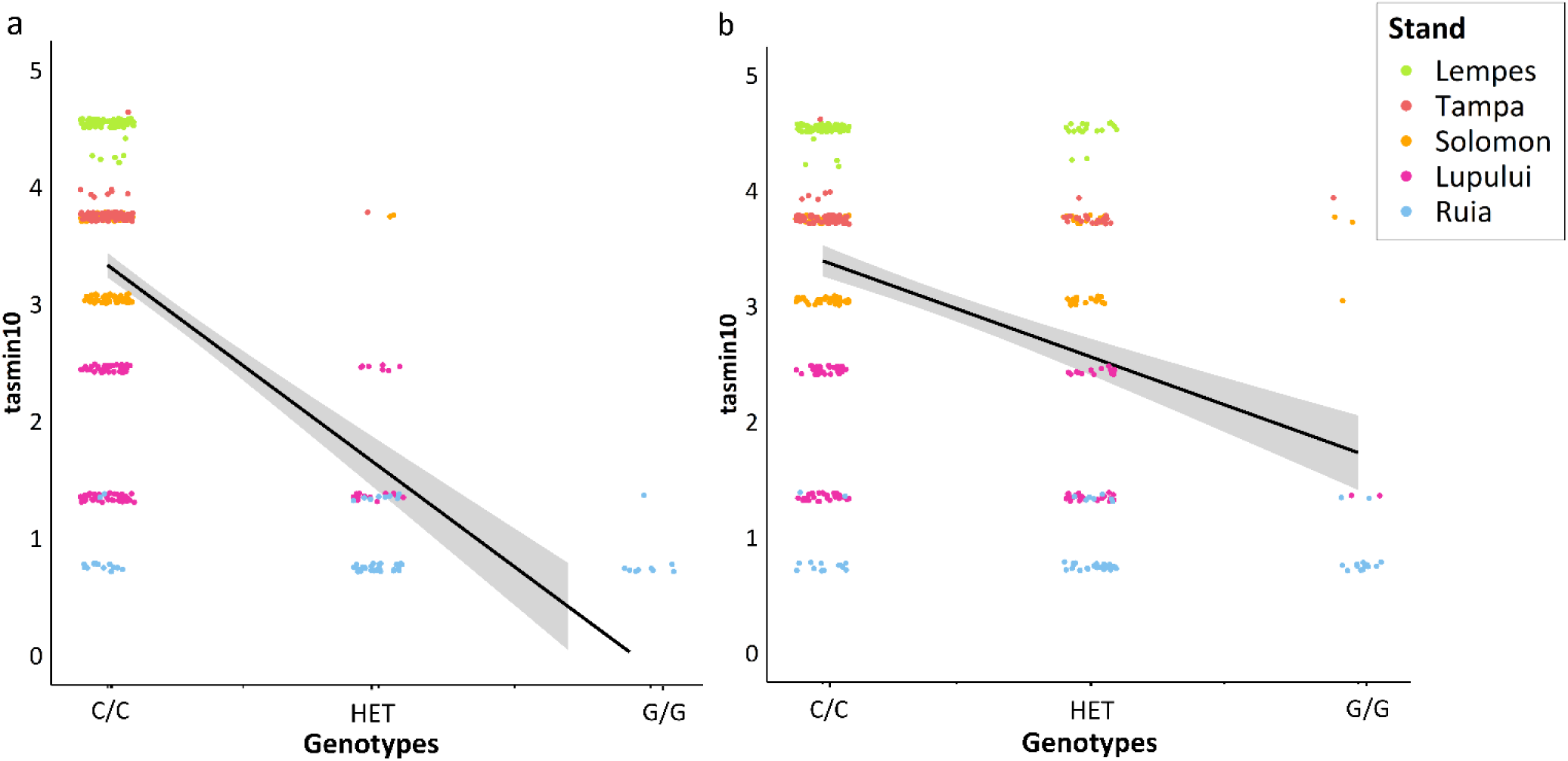
Correlation between minimum daily temperature in October (tasmin10) and observed genotypes at the significant SNP markers on chromosome 2 at ∼7.543 Mb (*Bhaga_2.g857,* with the underlying gene *polygalacturonase QRT3-like*) (**a**) and 7.787 Mb (*Bhaga_2.g887,* with the underlying gene protein *NRT1/ PTR FAMILY 5.4-like*) (**b**). The coefficients of determination R^2^ were 0.3468 (p ∼0) and 0.1646 (p ∼0).

## Discussion

### EAA results strongly correlate with temperature-based variables

This environmental association analysis aims to assess the adaptive potential to different environmental pressures along a steep environmental gradient in populations of European beech from five locations in the South-Eastern Romanian Carpathians. The altitudinal gradient is associated changes in precipitation and temperature (see supplemental Fig. S2). In total, 53 environmental variables and the principal components calculated based on these were used to perform environmental association analysis using LFMM (latent factor mixed models). We identified 446 SNP markers significantly associated with the first principal component (PC). These were overlapping with the SNP markers significantly associated with all environmental variables except precipitation accumulated during the growing season. The first PC was correlated with all temperature-based variables and elevation at |*r*| ∼0.989 to ∼0.997 and with all precipitation-based and Ellenberg-Quotient variables at |*r*| ∼0.945 to 0.950, except precipitation accumulated during the growing season. The 273 markers on chromosome 2 from ∼4.56 to ∼16.27 Mb, are overlapping or close to 22 genes. In total, we found ten genes with gene descriptions available based on our literature review (Table 1). Most of these genes are involved in gene regulation processes and enzyme catalyzation, and their impact is difficult to deduce (see supplemental Table S14). For some genes, functional studies are only available for homologs in rather distantly related species, such as humans (supplemental Table S14).

For two gene variants we found annotated genes potentially involved in local adaptation to specific environmental conditions and both genes were studied in plants. These gene variants were *Bhaga_2.g857* and *Bhaga_2.g883.* A high peak region on chromosome 2 from ∼4.56 to ∼16.27 Mb appeared in all results. This region was ∼3.47 Mb downstream from a high-confidence region for local adaptation identified by Lazic et al. (2024) on the same chromosome ∼0.79 Mb to 1.09 Mb.

### EEA peak region on chromosome 2 locates close to previously reported high-confidence region for local adaptation

Lazic et al. (2024) found the gene variant *Bhaga_2g.94* which was annotated with the gene *Callose synthase 1*. It was assumed that *Callose synthase 1* may be involved in winter dormancy and spring bud burst (Lazic et al. 2024). *Callose synthase 1* is described as being involved in callose deposition, during microspore development. Microspores are surrounded by a callose wall, this wall is then degraded and microspores are released (Tylewicz et al. 2018; Chu et al. 2024). We did not observe any significant SNP markers overlapping or in linkage disequilibrium (LD) with the high-confidence region from Lazic et al. (2024). However, we found a candidate gene based on our literature review, with very similar function to *Callose synthase 1* which was annotated for a marker on chromosome 2 at ∼7.543 Mb. This candidate gene is *polygalacturonase QRT3-like* (*Bhaga_2.g857*). The gene *polygalacturonase QRT3-like* plays a crucial role in pollen development in *Arabidopsis thaliana* L. (Rhee et al. 2003) and *Brassica rapa* L. (Chu et al. 2024). The gene function of *polygalacturonase QRT3-like* is similar to *Callose synthase 1* (Rhee et al. 2003; Tylewicz et al. 2018; Chu et al. 2024) and they are often studied together (Rhee et al. 2003; Chu et al. 2024). For further studies we recommend to not only consider the markers from this study, but also the region identified by Lazic et al. (2024). Our results provide statistical support for local adaptation on a small geographic scale in the face of gene flow along an environmental gradient. Lazic et al. (2024) studied local adaptation in 98 populations distributed from Southern Europe until Central and Eastern Europe.

### The candidate gene polygalacturonase QRT3-like

The gene variant *Bhaga_2.g857* corresponds to the underlying gene *polygalacturonase QRT3-like* (NCBI 2025). We found one SNP on chromosome 2 at ∼7.543 Mb located within the coding region of *Bhaga_2.g857 (polygalacturonase QRT3-like)* (supplemental Fig. S15). At this position of the coding region, a missense variant with moderate impact is located causing a codon change which leads to a change in the polar bonds of the amino acids (supplemental Table S16). The minor allele frequency at this marker decreased with increasing minimum daily temperature in October. The same significant correlation trend was observed for many temperature-based variables (see supplemental Fig. S11). The correlation with precipitation-based variables or other environmental variables like EQ, frost frequency change (fcf) and elevation was not significant. Our results suggest that this gene is involved in local adaptation to temperature.

Furthermore, a splice region variant of the coding region with low impact of this gene is located 70 bp upstream from the highly significant SNP marker (supplemental Fig. S15). The marker at the position of the splice region variant and the other missense variants are contained as markers in the data set, but are not significant. High LD is observed across the coding region of *Bhaga_2.g857 (polygalacturonase QRT3-like)*, but the LD is incomplete across the entire coding region of the gene (supplemental Fig. S15). It is possible that these markers are not exceeding the significance threshold because of insufficient power of the study (Sun et al. 2022). If very strict significance thresholds are chosen, only the coding region variants with the highest impact and at higher allele frequency in the population will be detected (Sun et al. 2022). In addition to the missense variants already mentioned, there are seven further missense variants between 21 bp upstream and 142 bp downstream of the marker (see supplemental Fig. S15).

The high-confidence region on chromosome 2 from ∼0.79 to 1.09 Mb from Lazic et al. (2024) was identified by three different methods for environmental association analysis (EAA). These three methods comprised LFMM from the LEA package (Frichot et al. 2013; Frichot and François 2015), Baypass (Gautier 2015) and WZA (Capblancq and Forester 2021). Additionally, Lazic et al. (2024) also measured the relative expression of *Callose synthase 1* in winter buds of eight beech trees. They observed that the relative expression level of *Callose synthase 1* was increased, but not differentially expressed, in the individuals carrying the alternative allele at the SNP marker identified as top LFMM hit on chromosome 2 at ∼0.898 Mb (Lazic et al. 2024). The individual trees carrying the homozygous genotype of the alternative allele at this SNP marker showed in a common garden experiment less days until bud burst (Lazic et al. 2024). Furthermore, a clear pattern of geographic distribution across the sampling area from Southern Europe to Central and Eastern Europe was observed (Lazic et al. 2024). The alternative allele was increased towards Central and Eastern Europe, in this region, the temperature of the coldest month was much lower (Lazic et al. 2024). To compare patterns of local adaptation at small and range-wide geographic scales, in future studies we will assess our candidate SNPs also at a larger geographic scale across the species distribution range.

### The candidate gene NRT1/PTR_FAMILY 5.4-like

For the gene variant *Bhaga_2.g883*, the annotated gene *NRT1/PTR_FAMILY 5.4-like* was found (NCBI 2025). The gene *NRT1/PTR_FAMILY 5.4-like* belongs to the NRT1/PTR family. This family was initially described as nitrate transporters, but recently their involvement in the transportation of phytohormones like auxin was discovered (Chiba et al. 2015; Watanabe et al. 2020). Many different members of this gene family are studied (Bai et al. 2013; Chiba et al. 2015; Watanabe et al. 2020), but *NRT1/PTR_FAMILY 5.4-like* was not studied in detail to our knowledge. But we can assume that this particular gene may also be involved in the transportation of phytohormones like auxin. The gene *NRT1/PTR_FAMILY* serves as a transporter for indole-3-butyric acid (IBA), which is a precursor of indole-acetic acid (IAA) (Watanabe et al. 2020). IAA is the major form of naturally available auxin (Watanabe et al. 2020; Chiba et al. 2015). At the position of the marker on chromosome 2 at ∼7.787 Mb, a missense variant with moderate impact was found in the coding region of *Bhaga_2.g883 (NRT1/PTR_FAMILY 5.4-like)* (supplemental Fig. S17). This missense variant causes a codon change which replaces a hydrophilic with a hydrophobic amino acid (supplemental Table S16). This change can increase the stability of the proteins (van Dijk et al. 2015) involved in auxin transport in the Ruia stands. Additionally, one splice region variant of the coding region of *Bhaga_2.g883 (NRT1/PTR_FAMILY 5.4-like)* was located 83 bp upstream from this significant marker (supplemental Fig. S17). This splice region variant was not recognized as significant in our analysis, which could also be due to the fact that it has less influence on the coding region. LD varies strongly across the gene coding region of *Bhaga_2.g883 (NRT1/PTR_FAMILY 5.4-like)* (supplemental Fig. S17). It seems like the splice region variant is not in strong LD with the marker and the rest of the coding region of this gene (supplemental Fig. S17). The missense variant in the coding region is in almost complete LD (∼1) with the marker (supplemental Fig. S17).

Figure 4B shows that the MAF at the marker on chromosome 2 at ∼ 7.787 Mb decreased with increasing minimum daily temperature in October. The same trend was observed for all temperature-based variables and also for elevation with a significance level of p≤0.05. The correlation coefficient for MAF at this marker and precipitation in October was ∼0.81 (p-value = 0.096), the correlation coefficients for the other precipitation-based variables except gsp were similar. Auxin activity is decreased during drought stress (Popko et al. 2009). A decrease in free auxin in the cambial zone of poplar was observed in acclimation to drought stress caused by osmotic stress due to changes in water potential in the environment (Popko et al. 2009). In general, high auxin activity is correlated with high growth rates in trees (Popko et al. 2009). Since this particular gene was not studied, it is difficult to draw direct conclusions about the exact involvement in local adaptation, but we would assume that auxin is decreased in the shoot apical meristems of the stands with high drought stress. The allele frequency at the SNP candidate marker on chromosome 2 at ∼7.787 Mb is decreased in the stands Lempes, Tampa and Solomon with higher drought stress (see Fig. 4b). Only in Ruia, the allele frequency at the SNP candidate is increased (see Fig. 4b).

Both gene variants seem to be involved in local adaptation. The LD between the two SNP markers on chromosome 2 at ∼ 7.543 and ∼ 7.787 is 0.6711. The LD across the entire region from ∼4.56 to ∼16.27 Mb on chromosome 2 varies (Fig. 3). It seems like temperature-based variables are shaping local adaptation much stronger then precipitation-based variables, especially precipitation accumulated over the growing season does not seem to cause strong patterns of local adaptation, at least in these stands.

The coefficients of determination of the linear regression between SNP genotypes and minimum daily temperature in October on chromosome 2 at ∼ 7.543 Mb and at ∼7.787 Mb are 0.3468 and 0.1646, respectively (Fig. 5). For the SNP maker at ∼ 7.543 Mb, the homozygous G/G genotype only occurs in the high-elevation population Ruia, while the heterozygous genotype C/G is mostly restricted to the high elevation populations Ruia and Lupului. On the other hand, the C/C genotype is present in similar frequencies in all populations. This pattern suggests strong environmental selection. This trend is not as pronounced for the marker at ∼7.787 Mb on chromosome 2 (Fig. 5).

In a genome-wide association study (GWAS) conducted on the same genotyping data set as in this study, we observed significant associations between 101 SNPs located on chromosome 10 and stomatal density as shown in Tost et al. (2025). These 101 markers are located on chromosome 10 in a region spreading from ∼4.9 to 13.67 Mb and also appear as one region (Tost et al. 2025). Five genes were located in this region that may play a role in controlling stomatal density (Tost et al. 2025). All markers within this region exhibit similar allele frequencies, which are correlated with stomatal density and the altitudinal gradient of the stands (see Tost et al. 2025). For the trait carbon isotope composition δ^13^C measured in 2020 and 2022, we observed also a peak on chromosome 10, which was much smaller and only resulted in 2020 in two significant SNP marker associations (Tost et al. 2025). Other significant markers associated with leaf nitrogen content or C/N ratio were also located on chromosome 2 but at ∼30.26 Mb and ∼31.804 Mb, ∼14 Mb downstream of the region in this study (Tost et al. 2025). For the significant SNP marker on chromosome 2 at ∼31.804 Mb associated with C/N ratio measured in 2021, the gene variant *Bhaga_2.g3457* was found based on a literature review. *Bhaga_2.g3457* was annotated as *abscisic-aldehyde oxidase-like* (NCBI 2024). The gene *abscisic-aldehyde oxidase-like* is responsible for oxidation of abscisic aldehyde, which is the last step of abscisic acid (ABA) biosynthesis (Seo et al. 2000). Popko et al. (2009) observed under drought stress auxin and abscisic acid (ABA) interact to regulate plant water status in poplar. The markers identified in our EAA and the GWAS do not overlap, possibly due to insufficient power of the analyses and multiple traits under environmental selection.

### Permutation testing

For the significance thresholds based on the BH correction, an average of 39 false-positive results were observed for the ten different permutations with ten replications (supplementary Table S6). We identified 446 SNP markers significantly associated with the first principal component (PC), which are considered to be putatively true positive observations. Based on our permutation tests, we calculated a false positive rate of ∼0.09. This false-positive rate of ∼0.09 is relatively low, so we did not adjust the significance thresholds.

## Conclusions

In conclusion, our research confirms that environmental association analysis (EAA) can identify environmental variables, which contribute to local adaptation. We observed that mainly temperature-based variables were detected with EAA and showed the strongest correlation with minor allele frequency (MAF) at significant SNP markers from the EAA. Consequently, the adaptive potential to temperature along a steep environmental gradient in the European beech populations was relatively high. A region for local adaptation on chromosome 2 from ∼4.56 to ∼16.27 Mb was identified based on this study and associated with all environmental variables except precipitation accumulated over the growing season. It seems like temperature-based variables shaped local adaptation. The candidate SNP markers on chromosome 2 at ∼7.543 (*Bhaga_2.g857)* and at ∼7.787 (*Bhaga_2.g883*) were annotated with the genes *polygalacturonase QRT3-like* and *NRT1/PTR_FAMILY 5.4-like*. The gene *polygalacturonase QRT3-like* plays a role in pollen development in Arabidopsis. The gene *NRT1/PTR_FAMILY 5.4-like* belongs to the NRT1/PTR family, which is involved in the transportation of phytohormones like auxin. The MAF observed at these candidate SNP markers is increased especially in Ruia, the population at highest altitude. Both candidate SNP markers are annotated at a part of the coding region of both genes where a codon change replaces a hydrophilic with a hydrophobic amino acid. This codon change may increase the stability of proteins encoded by genes involved in auxin transport and may be involved in pollen development in the Ruia stands. This is very speculative, because in order to substantiate these assumptions, expression levels of the genes need to measured.

## Availability of data and materials

Raw data is publicly available at figshare: https://figshare.com/articles/dataset/VCF_file_with_SPET_sequencing_data_of_DroughtMarkers_project/28748924 and was published by Tost et al. (2025). Our analysis code is available publicly on github: https://github.com/milaleonie/DroughtMarkers.

## Acknowledgements

We would like to thank all people involved in this project. We are thanking for help in the lab our lab technician Alexandra Dolynska and bachelor and master students Nikolas Heeger, Luca Schwendemann, Merle Kleikamp, Timo Unbehau, and Jonas Daniel. Furthermore, we are thanking Dr. Mehdi Ben Targem, Mihnea-Ioan-Cezar Ciocîrlan, Jürgen Scheuring and Jonah Fels for help with sampling in Romania.

## Funding

The used data comes from the drought markers project (Reference numbers: 2218WK43B4, 2218WK43A4). This work was financially supported by the Federal Ministry of Food and Agriculture – FNR-Waldklimafonds. The drought markers project is a collaboration between the University of Goettingen (Germany), the HAWK (Germany), the NW-FVA (Germany) and Transilvania University of Brasov (Romania). We utilized the computational resources of the University of Goettingen’s GWDG. Open Access funding enabled and organized by Projekt DEAL.

## Conflict of interest

The authors declare that there is no conflict of interest.

## Notes

### Competing Interest Statement

The authors have declared no competing interest.

