## Supplemental Files for "Assessing the adaptative potential to temperature and precipitation along a steep environmental gradient in populations of European beech"

**Supplementary files**

**S1:** Distribution of diameter at breast height (DBH) measurements across stands.

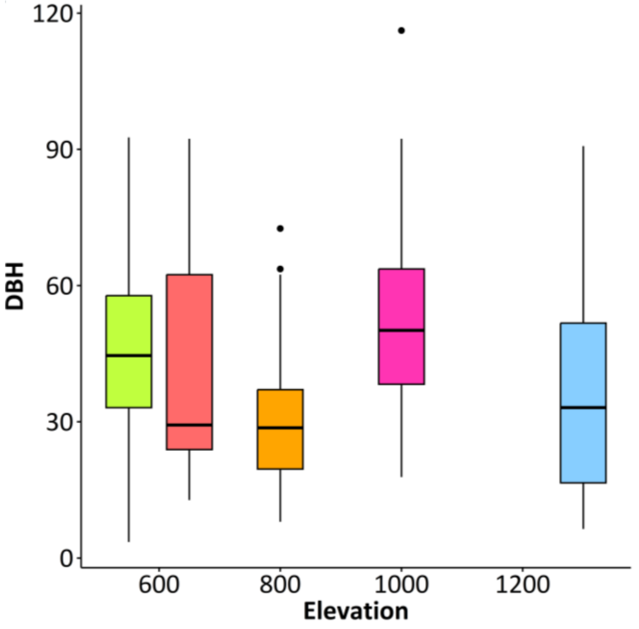


**S2:** Distribution of environmental variables across stands with the precipitation based variables CHE_pr_12, CHE_pr_11, CHE_pr_10, CHE_pr_09, CHE_pr_08, CHE_pr_07, CHE_pr_06, CHE_pr_05, CHE_pr_04, CHE_pr_03, CHE_pr_02 and CHE_pr_01 (precipitation in December, November, October, September, August, July, June, May, April, March, February and January) and temperature-based variables tas12, tas11, tas10, tas09 tas08, tas07, tas06, tas05, tas04, tas03, tas02 and tas01 (temperature in December, November, October, September, August, July, June, May, April, March, February and January).

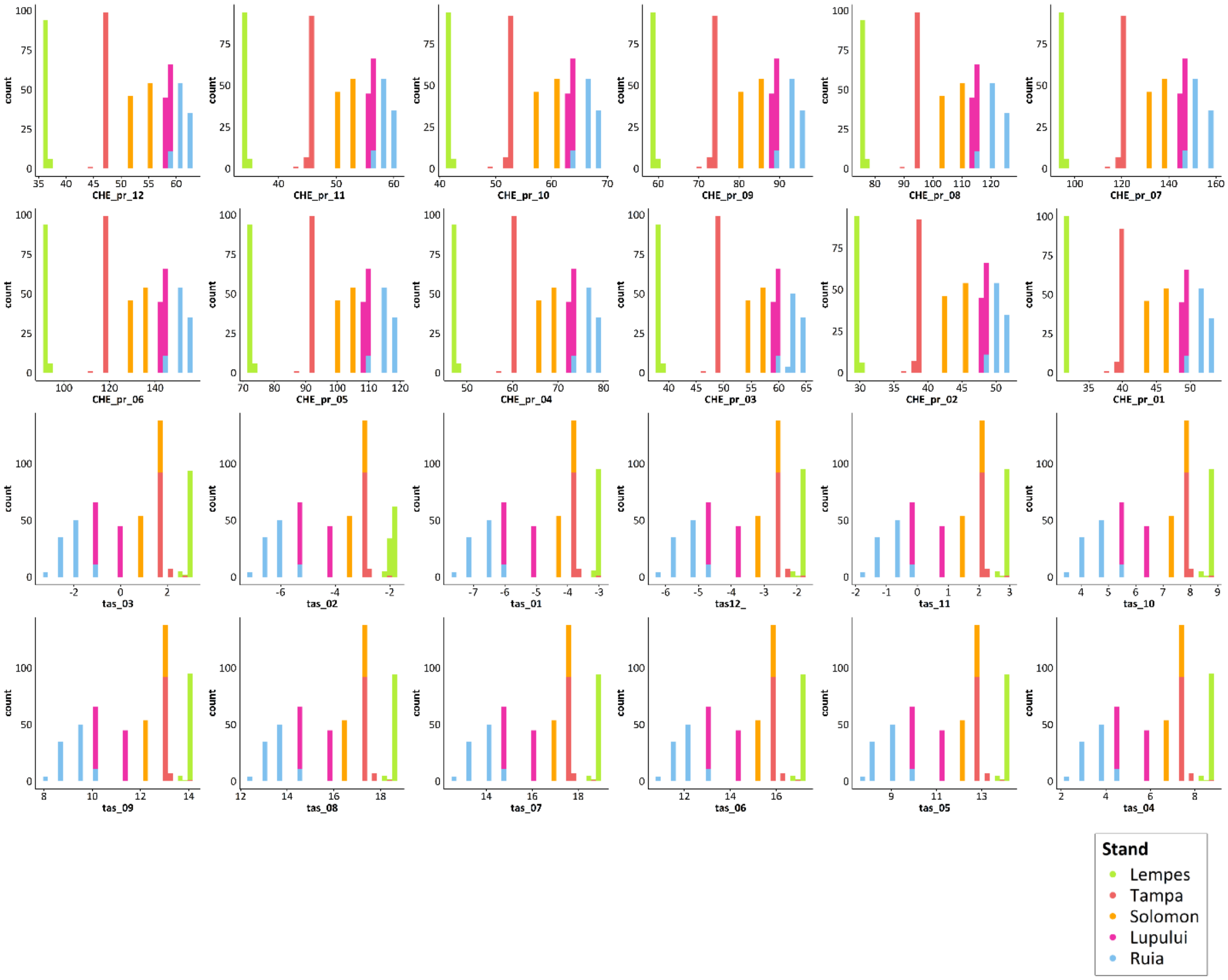


**S3:** Correlations of all environmental variables and their correlation coefficients. All correlations are significant at p~0***.

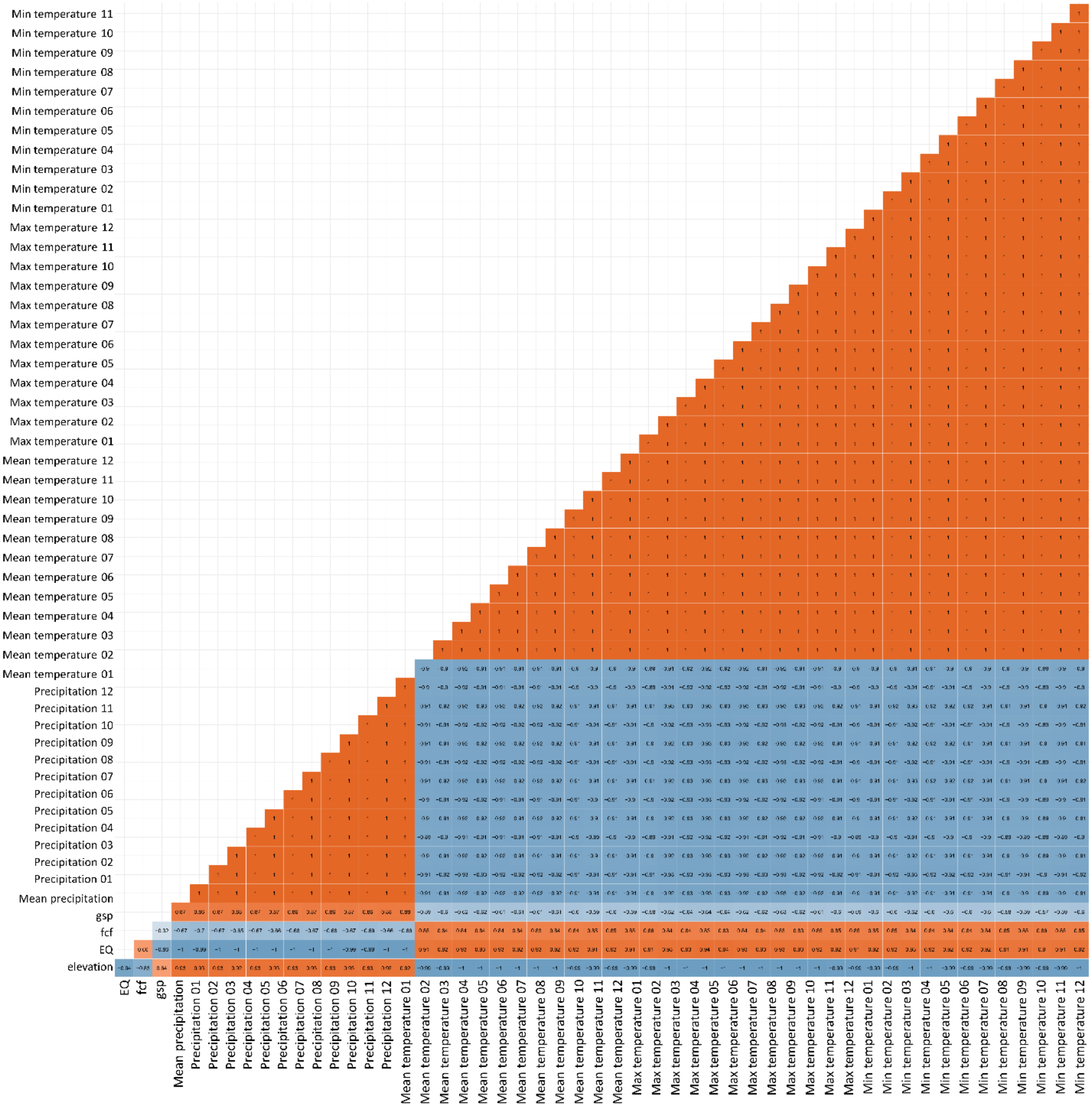


**S4:** Principal component analysis (PCA) based on all 53 environmental variables with principal component 1 plotted against principal component 2 (A) and the eigenvalues of the other principal components.

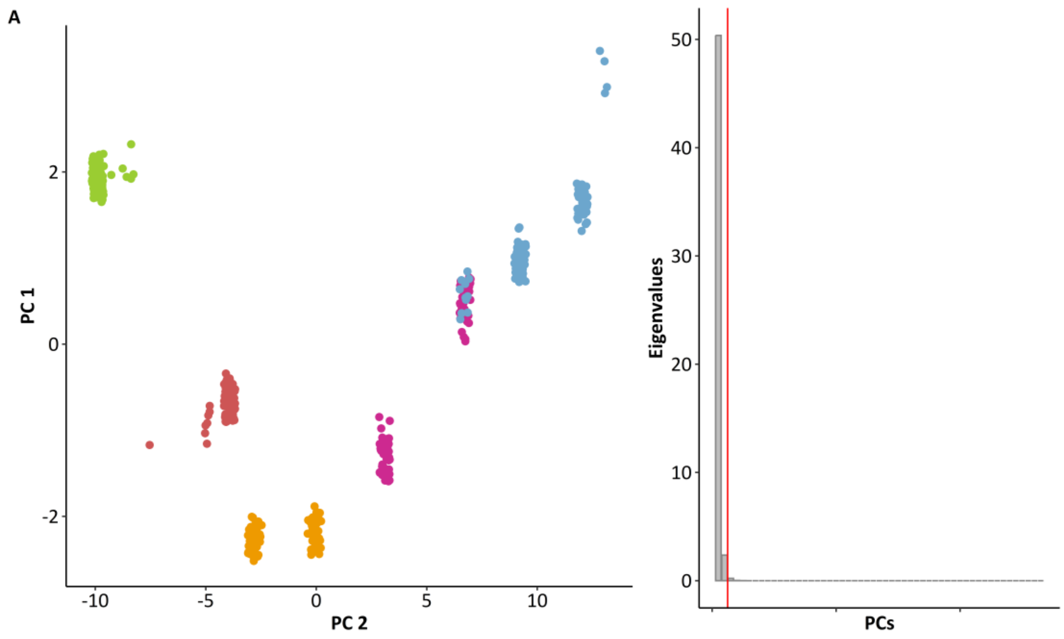


**S5:** Distribution of missingness (marker coverage) across samples before lfmm imputation with cut-off threshold at a missingness of 0.05 (red) which corresponds to a marker coverage of 0.95.

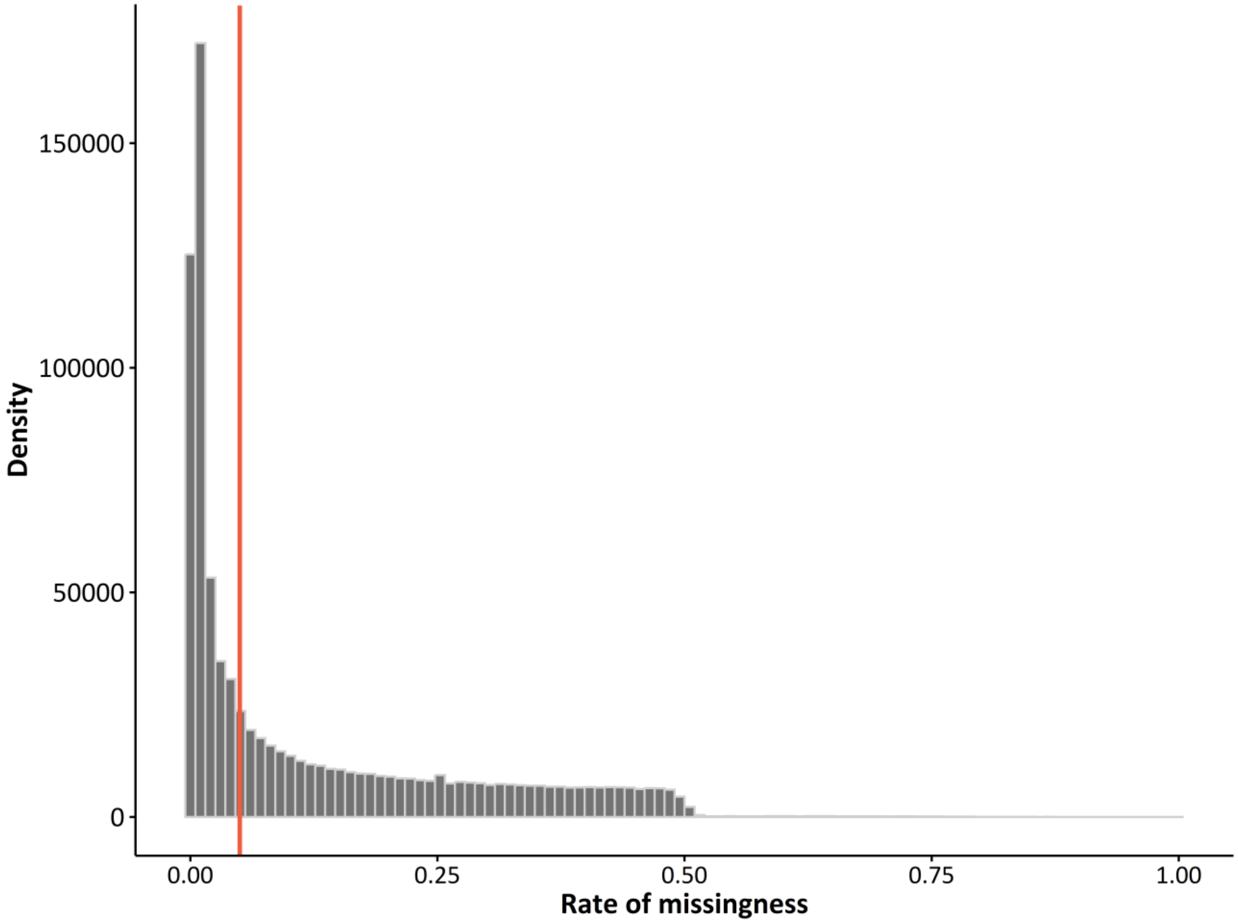

**S6:** Randomization scheme and results from permutations (perm.) with number of false positive observations per replications. In the randomization scheme the order of the stands Ruia (Ru), Lupului (Lu), Solomon (So), Tampa (Ta) and Lempes (Le) was changed. The table also shows the initial order of the stands and the results from the not permuted analysis.

| **Real** | **Initial order of stands** | | | | | **Number of observations** | | | | | | | | | | |
|---|---|---|---|---|---|---|---|---|---|---|---|---|---|---|---|---|
| | Ru | Lu | So | Ta | Le | 446 | | | | | | | | | | |
| **Perm.** | **Randomization scheme** | | | | | **Number of false positive observations per replication** | | | | | | | | | | **μ** |
| Run 1 | Lu | Le | Ta | Ru | So | 249 | 255 | 265 | 265 | 265 | 261 | 263 | 259 | 247 | 249 | 257.61 |
| Run 2 | So | Ru | Le | Ta | Lu | 198 | 193 | 203 | 207 | 207 | 193 | 201 | 198 | 219 | 204 | 202.04 |
| Run 3 | Ta | Ru | So | Lu | Le | 273 | 207 | 260 | 264 | 247 | 259 | 265 | 265 | 268 | 268 | 256.10 |
| Run 4 | Le | Ru | Ta | Lu | So | 225 | 231 | 235 | 243 | 243 | 231 | 235 | 216 | 221 | 224 | 230.08 |
| Run 5 | Ta | Lu | So | Ru | Le | 7 | 6 | 5 | 7 | 7 | 7 | 8 | 6 | 6 | 7 | 6.50 |
| Run 6 | Lu | Ta | Le | Ru | So | 12 | 13 | 13 | 11 | 11 | 12 | 12 | 13 | 11 | 12 | 11.95 |
| Run 7 | Le | Lu | Ta | Ru | So | 8 | 9 | 9 | 9 | 9 | 9 | 9 | 12 | 8 | 9 | 9.00 |
| Run 8 | So | Lu | Ta | Ru | Le | 8 | 2 | 2 | 2 | 2 | 3 | 2 | 3 | 4 | 2 | 2.47 |
| Run 9 | Ru | Le | So | Lu | Ta | 297 | 309 | 305 | 309 | 305 | 305 | 298 | 297 | 305 | 311 | 304.02 |
| Run 10 | Ru | Ta | Lu | Le | So | 6 | 6 | 7 | 6 | 5 | 3 | 3 | 6 | 4 | 5 | 4.70 |
| | | | | | | | | | | | | | | | | 38.72 |

**S7:** Analysis of molecular variance (AMOVA) to assess the variation between stands and within stands with the poppr R package (Kamvar et al. 2013) expressed in %.

| **Source** | **Df** | **Sum Sq** | **Mean Sq** | **Sigma** | **%** |
|---|---|---|---|---|---|
| **Variations between samples** | 4 | 410787.5 | 102696.9 | 786.344 | 3.11 |
| **Variations within samples** | 492 | 12071098 | 24534.75 | 24534.75 | 96.89 |
| **Total variations** | 496 | 12481885 | 25165.09 | 25321.1 | 100 |

**S8:** Pairwise $F_{ST}$ matrix calculated based on all SNP markers with the StAMPP R package (Pembleton et al. 2013).

| | **Ruia** | **Lupului** | **Solomon** | **Tampa** | **Lempes** |
|---|---|---|---|---|---|
| **Ruia** | NA | NA | NA | NA | NA |
| **Lupului** | 0.000459 | NA | NA | NA | NA |
| **Solomon** | -0.000028 | -0.003601 | NA | NA | NA |
| **Tampa** | 0.000131 | -0.004026 | -0.003885 | NA | NA |
| **Lempes** | -0.003936 | 0.000546 | -0.000036 | 0.000178 | NA |

**S9:** Manhattan plots based on LFMM results for environmental PC 2 (a), PC 3 (b), PC 4 (c), PC 5 (d), PC 6 (e), PC 7 (f), PC 8 (g) with Benjamini-Hochberg (BH) corrected p-values on the –log10.

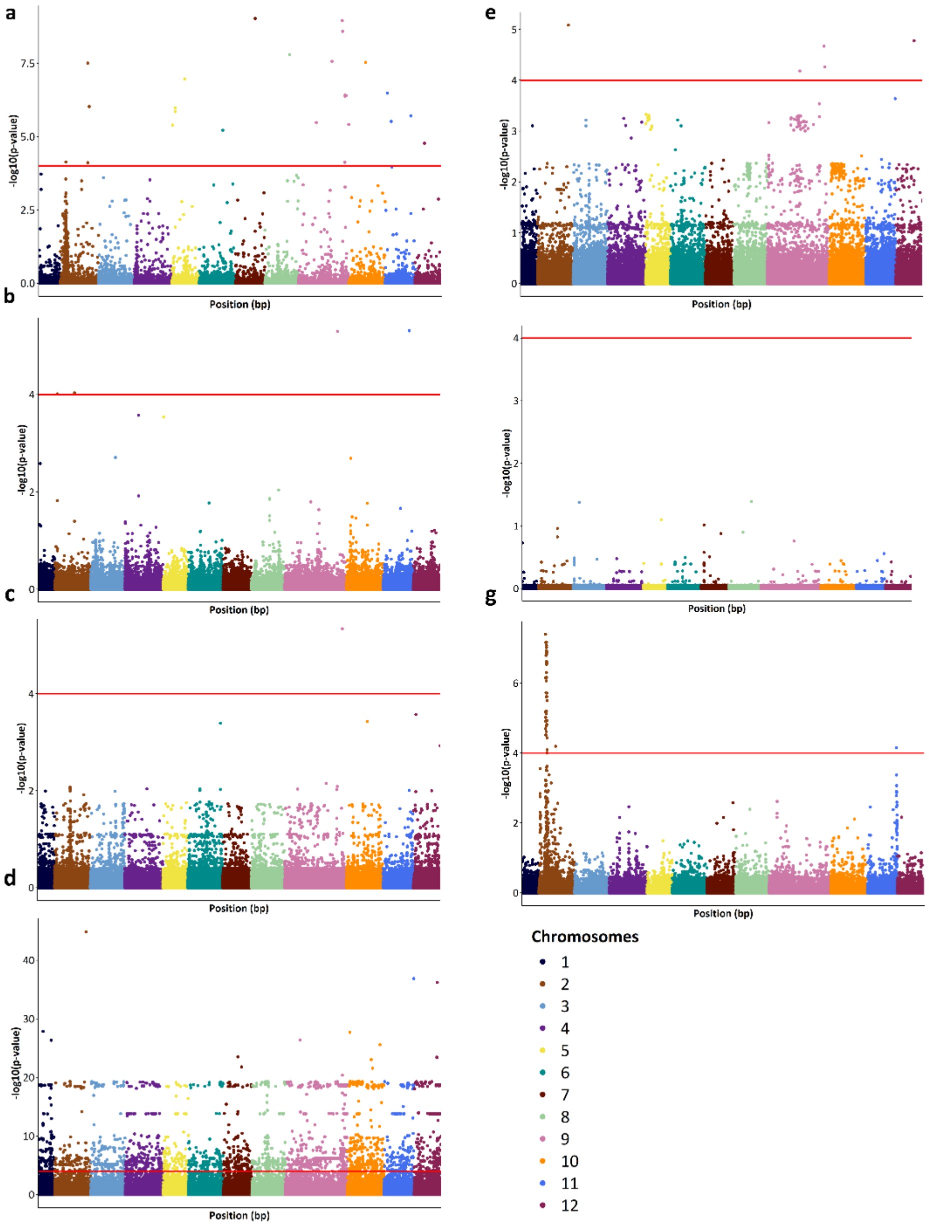


**S10:** Manhattan plots based on LFMM results for mean precipitation (**a**), daily mean temperature (**b**), daily maximum (**c**), daily minimum temperature in July (**d**), elevation (**e**), EQ (**f**) and precipitation accumulated over the growing season (gsp) (**f**) with Benjamini-Hochberg (BH) corrected p-values on the –log10 .

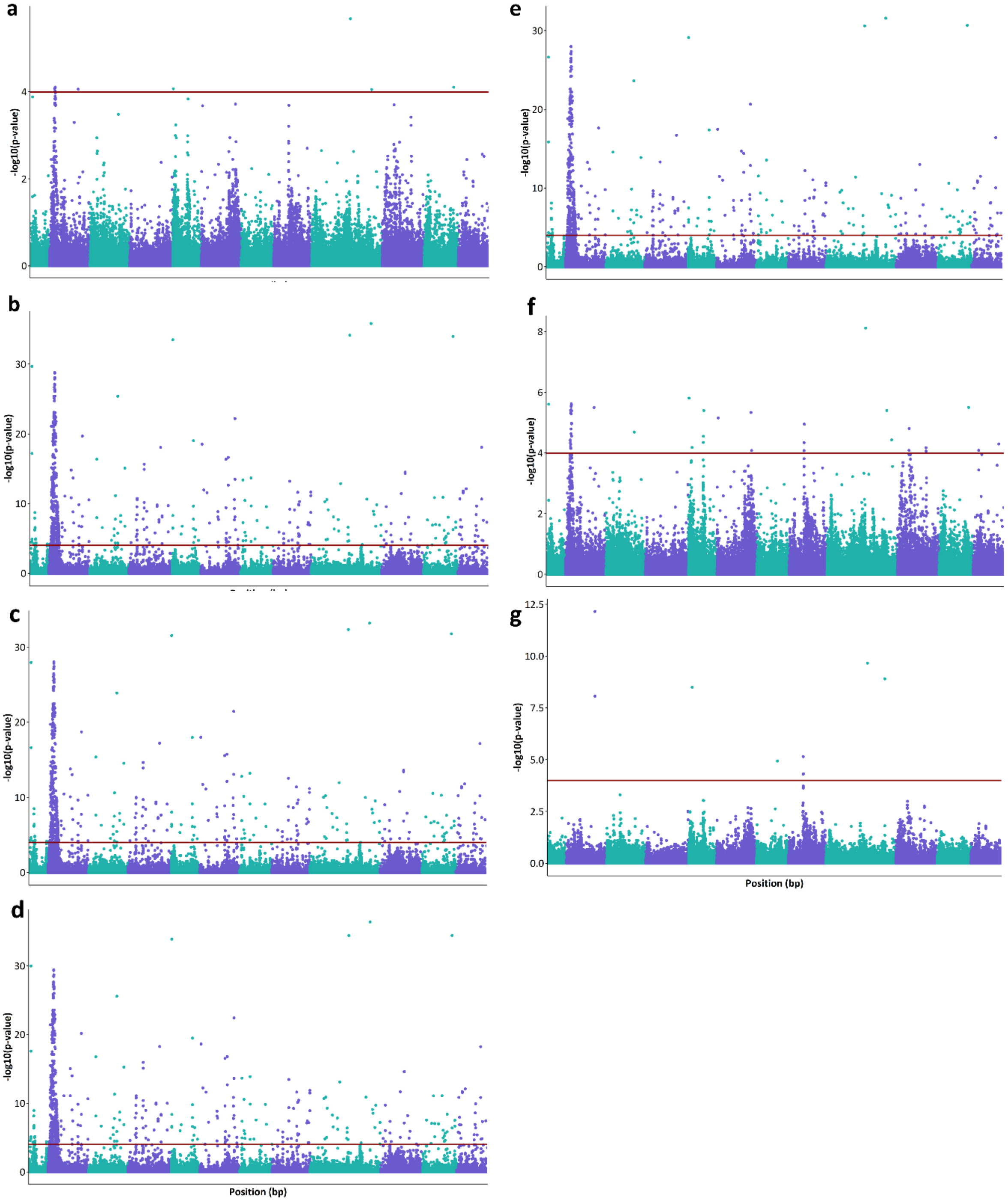


**S11:** Correlation between with different environmental variables tas01, tas06, tas09 (maximum daily temperature in January, June, September) and tasMAX01, tasMAX06, tasMAX09 (daily temperature in January, June, September) at the significant marker on chromosome 2 at ~7.543 (*Bhaga_2.g857)* and at ~7.787 (*Bhaga_2.g883)*. These markers were annotated with gene variant *Bhaga_2.g857*, with the underlying gene *polygalacturonase QRT3-like* and with gene variant *Bhaga_2.g883*, with the underlying gene protein *NRT1/ PTR FAMILY 5.4-like*.

### Marker on chromosome 2 at ~ 7.543 Mb

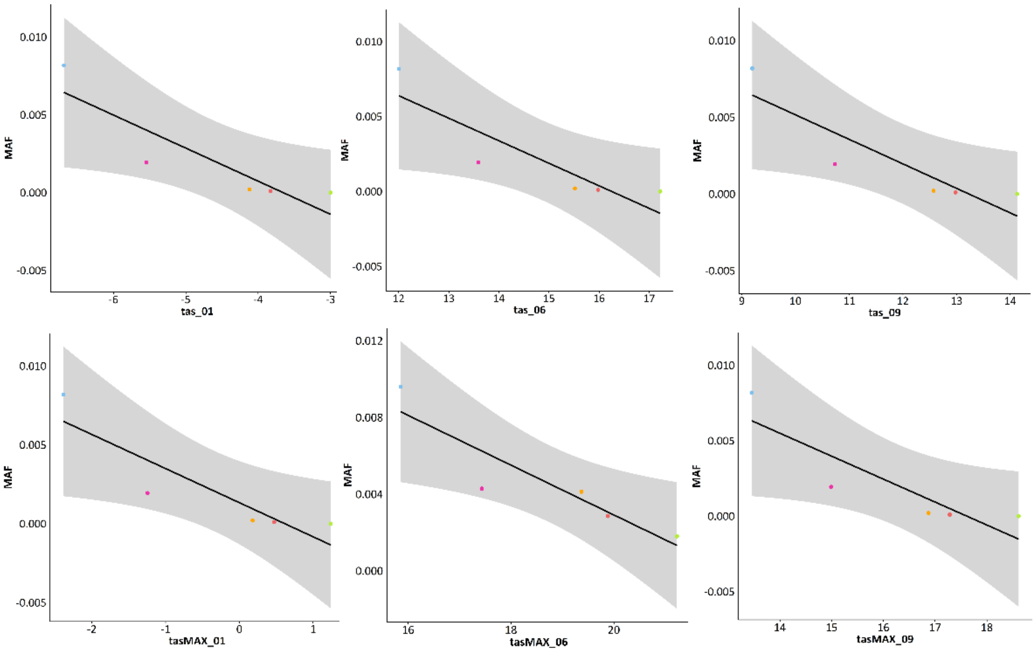


### Marker on chromosome 2 at ~ 7.787 Mb

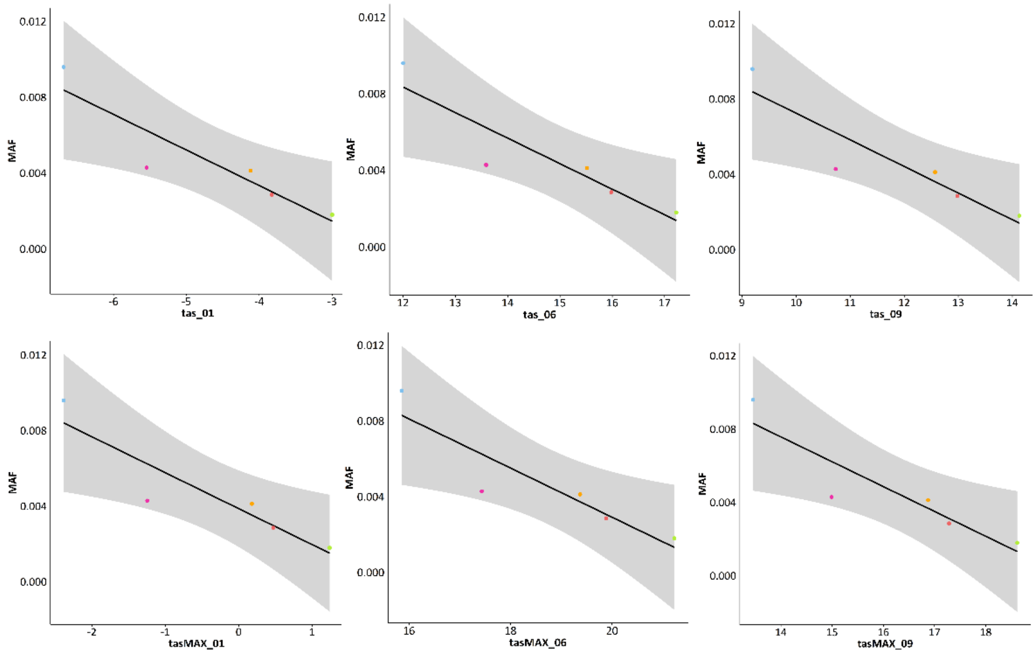


**S12:** Minor allele frequency (MAF) observed in the different stands at the significant markers ($p \leq 0.0001$) on chromosome 2 associated with the different environmental variables tasMAX01, tasMAX02, tasMAX06, tasMAX09, tasMAX10, tasMAX11, tasMAX12 (maximum daily temperature in January, February, June, September, October, November and December), tasmin01, tasmin02, tasmin07, tasmin11, tasmin12 (minimum daily temperature in January, February, July, November, December) and elevation and their mean.

| Chr | Marker | Stand | MAF | Env. variable | Mean | Chr | Marker | Stand | MAF | Env. variable | Mean |
|---|---|---|---|---|---|---|---|---|---|---|---|
| 2 | 7542764 | Lempes | 0.0000 | tasMAX01 | 1.239 | 2 | 7787374 | Lempes | 0.0900 | tas07 | 18.89 |
| 2 | 7542764 | Tampa | 0.0051 | tasMAX01 | 0.472 | 2 | 7787374 | Tampa | 0.1414 | tas07 | 17.583 |

| | | | | | | | | | | | |
|---|---|---|---|---|---|---|---|---|---|---|---|
| 2 | 7542764 | Solomon | 0.0556 | tasMAX01 | 0.18 | 2 | 7787374 | Solomon | 0.2475 | tas07 | 17.172 |
| 2 | 7542764 | Lupului | 0.0960 | tasMAX01 | -1.245 | 2 | 7787374 | Lupului | 0.2121 | tas07 | 15.335 |
| 2 | 7542764 | Ruia | 0.4500 | tasMAX01 | -2.384 | 2 | 7787374 | Ruia | 0.5200 | tas07 | 13.756 |
| 2 | 7542764 | Lempes | 0.0000 | tasMAX02 | 2.628 | 2 | 7787374 | Lempes | 0.0900 | tasmin11 | -0.467 |
| 2 | 7542764 | Tampa | 0.0051 | tasMAX02 | 1.573 | 2 | 7787374 | Tampa | 0.1414 | tasmin11 | -1.32 |
| 2 | 7542764 | Solomon | 0.0556 | tasMAX02 | 1.126 | 2 | 7787374 | Solomon | 0.2475 | tasmin11 | -1.62 |
| 2 | 7542764 | Lupului | 0.0960 | tasMAX02 | -0.5 | 2 | 7787374 | Lupului | 0.2121 | tasmin11 | -3.1 |
| 2 | 7542764 | Ruia | 0.4500 | tasMAX02 | -1.812 | 2 | 7787374 | Ruia | 0.5200 | tasmin11 | -4.284 |
| 2 | 7542764 | Lempes | 0.0000 | tasMAX06 | 21.228 | 2 | 7787374 | Lempes | 0.0900 | tasmin12 | -5.066 |
| 2 | 7542764 | Tampa | 0.0051 | tasMAX06 | 19.883 | 2 | 7787374 | Tampa | 0.1414 | tasmin12 | -5.921 |
| 2 | 7542764 | Solomon | 0.0556 | tasMAX06 | 19.372 | 2 | 7787374 | Solomon | 0.2475 | tasmin12 | -6.22 |
| 2 | 7542764 | Lupului | 0.0960 | tasMAX06 | 17.435 | 2 | 7787374 | Lupului | 0.2121 | tasmin12 | -7.645 |
| 2 | 7542764 | Ruia | 0.4500 | tasMAX06 | 15.856 | 2 | 7787374 | Ruia | 0.5200 | tasmin12 | -8.745 |
| 2 | 7542764 | Lempes | 0.0000 | tasMAX09 | 18.614 | 2 | 7787374 | Lempes | 0.0900 | tas01 | -2.999 |
| 2 | 7542764 | Tampa | 0.0051 | tasMAX09 | 17.283 | 2 | 7787374 | Tampa | 0.1414 | tas01 | -3.828 |
| 2 | 7542764 | Solomon | 0.0556 | tasMAX09 | 16.872 | 2 | 7787374 | Solomon | 0.2475 | tas01 | -4.12 |
| 2 | 7542764 | Lupului | 0.0960 | tasMAX09 | 14.99 | 2 | 7787374 | Lupului | 0.2121 | tas01 | -5.545 |
| 2 | 7542764 | Ruia | 0.4500 | tasMAX09 | 13.456 | 2 | 7787374 | Ruia | 0.5200 | tas01 | -6.684 |
| 2 | 7542764 | Lempes | 0.0000 | tasMAX10 | 13.527 | 2 | 7787374 | Lempes | 0.0900 | tas02 | -1.904 |
| 2 | 7542764 | Tampa | 0.0051 | tasMAX10 | 12.473 | 2 | 7787374 | Tampa | 0.1414 | tas02 | -2.919 |
| 2 | 7542764 | Solomon | 0.0556 | tasMAX10 | 12.026 | 2 | 7787374 | Solomon | 0.2475 | tas02 | -3.274 |
| 2 | 7542764 | Lupului | 0.0960 | tasMAX10 | 10.345 | 2 | 7787374 | Lupului | 0.2121 | tas02 | -4.855 |
| 2 | 7542764 | Ruia | 0.4500 | tasMAX10 | 9.023 | 2 | 7787374 | Ruia | 0.5200 | tas02 | -6.177 |
| 2 | 7542764 | Lempes | 0.0000 | tasMAX11 | 7.133 | 2 | 7787374 | Lempes | 0.0900 | tas03 | 3.028 |
| 2 | 7542764 | Tampa | 0.0051 | tasMAX11 | 6.272 | 2 | 7787374 | Tampa | 0.1414 | tas03 | 1.782 |
| 2 | 7542764 | Solomon | 0.0556 | tasMAX11 | 5.88 | 2 | 7787374 | Solomon | 0.2475 | tas03 | 1.272 |
| 2 | 7542764 | Lupului | 0.0960 | tasMAX11 | 4.4 | 2 | 7787374 | Lupului | 0.2121 | tas03 | -0.61 |
| 2 | 7542764 | Ruia | 0.4500 | tasMAX11 | 3.162 | 2 | 7787374 | Ruia | 0.5200 | tas03 | -2.105 |
| 2 | 7542764 | Lempes | 0.0000 | tasMAX12 | 2.233 | 2 | 7787374 | Lempes | 0.0900 | tas04 | 8.822 |
| 2 | 7542764 | Tampa | 0.0051 | tasMAX12 | 1.472 | 2 | 7787374 | Tampa | 0.1414 | tas04 | 7.483 |
| 2 | 7542764 | Solomon | 0.0556 | tasMAX12 | 1.18 | 2 | 7787374 | Solomon | 0.2475 | tas04 | 7.018 |
| 2 | 7542764 | Lupului | 0.0960 | tasMAX12 | -0.145 | 2 | 7787374 | Lupului | 0.2121 | tas04 | 5.09 |
| 2 | 7542764 | Ruia | 0.4500 | tasMAX12 | -1.249 | 2 | 7787374 | Ruia | 0.5200 | tas04 | 3.502 |
| 2 | 7542764 | Lempes | 0.0000 | tasmin01 | -6.899 | 2 | 7787374 | Lempes | 0.0900 | tas05 | 14.122 |
| 2 | 7542764 | Tampa | 0.0051 | tasmin01 | -7.728 | 2 | 7787374 | Tampa | 0.1414 | tas05 | 12.79 |
| 2 | 7542764 | Solomon | 0.0556 | tasmin01 | -8.074 | 2 | 7787374 | Solomon | 0.2475 | tas05 | 12.372 |
| 2 | 7542764 | Lupului | 0.0960 | tasmin01 | -9.445 | 2 | 7787374 | Lupului | 0.2121 | tas05 | 10.435 |
| 2 | 7542764 | Ruia | 0.4500 | tasmin01 | - | 2 | 7787374 | Ruia | 0.5200 | tas05 | 8.856 |
| 2 | 7542764 | Lempes | 0.0000 | tasmin02 | -6.567 | 2 | 7787374 | Lempes | 0.0900 | tas06 | 17.222 |
| 2 | 7542764 | Tampa | 0.0051 | tasmin02 | -7.427 | 2 | 7787374 | Tampa | 0.1414 | tas06 | 15.982 |
| 2 | 7542764 | Solomon | 0.0556 | tasmin02 | -7.874 | 2 | 7787374 | Solomon | 0.2475 | tas06 | 15.518 |
| 2 | 7542764 | Lupului | 0.0960 | tasmin02 | -9.4 | 2 | 7787374 | Lupului | 0.2121 | tas06 | 13.59 |
| 2 | 7542764 | Ruia | 0.4500 | tasmin02 | - | 2 | 7787374 | Ruia | 0.5200 | tas06 | 12.006 |
| 2 | 7542764 | Lempes | 0.0000 | tasmin07 | 13.628 | 2 | 7787374 | Lempes | 0.0900 | elevation | 550.82 |
| 2 | 7542764 | Tampa | 0.0051 | tasmin07 | 12.483 | 2 | 7787374 | Tampa | 0.1414 | elevation | 766.86 |
| 2 | 7542764 | Solomon | 0.0556 | tasmin07 | 12.072 | 2 | 7787374 | Solomon | 0.2475 | elevation | 852.12 |
| 2 | 7542764 | Lupului | 0.0960 | tasmin07 | 10.235 | 2 | 7787374 | Lupului | 0.2121 | elevation | 1144.46 |
| 2 | 7542764 | Ruia | 0.4500 | tasmin07 | 8.695 | 2 | 7787374 | Ruia | 0.5200 | elevation | 1376.22 |

**S13:** Correlation between tasmin10 (minimum daily temperature in October) and the minor allele frequency (MAF) of the significant marker (p ≤0.0001) on chromosome 2 at **a**) ~ 10.07112 Mb at-0.8566 (p-value = 0.0638), **b**) ~10.07115 Mb at -0.8481 (p-value = 0.0694), **c**) ~ 10.526 Mb at -0.9201 (p-value = 0.0268), **d**) ~ 10.533 Mb at -0.9119 (p-value = 0.0309), **e**) ~ 11.929 Mb at -0.9055 (p-value = 0.0344), **f**) ~ 11.989 Mb at -0.9012 (p-value = 0.0367), **g**) ~ 14.682 Mb at -0.9236 (p-value = 0.0250), **h**) ~ 14.823 Mb at -0.8929 (p-value = 0.0414), **i**) ~ 15.895 Mb at -0.9148 (p-value = 0.0295), **j**) ~ 16.130 Mb at -0.8745 (p-value = 0.0523).

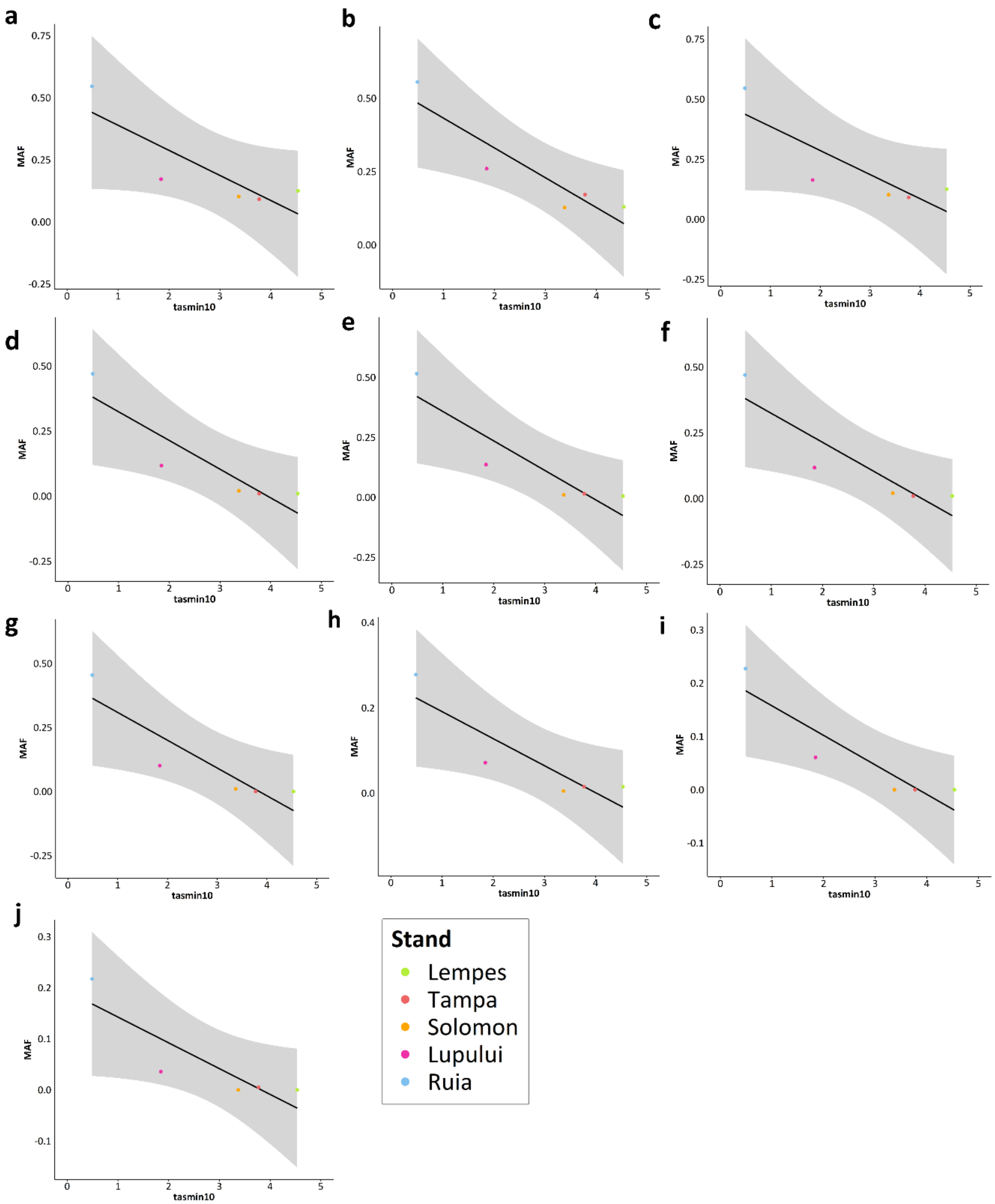


**S14:** List of gene descriptions close to markers associated with environmental PC1

| Gene variant name | Gene | Gene description based on the literature review |
|---|---|---|
| **Bhaga_2.g1166** | **transcription elongation factor SPT6 homolog** | **Transcription elongation factor SPT6 homolog** is an essential and conserved transcription factor which directly binds to and co-localizes with RNAPII at sites of transcription (Solano et al. 2017). |

| | | |
|---|---|---|
| **Bhaga_2.g1303** | **NAD-dependent malic enzyme 62 kDa isoform, mitochondrial** | Malic enzymes catalyze the oxidative decarboxylation of malate to pyruvate and $CO_2$ (Xu et al. 1999). |
| **Bhaga_2.g1308** | **ras-related protein RABH1b** | **ras-related protein** RABH1b was observed to be upregulated under salinity (Manivannan et al. 2016). In general, Ras-proteins play an essential role in signal transduction pathways that control cellular processes (Talajić et al. 2024). |
| **Bhaga_2.g1609** | **spindle and kinetochore-associated protein 1 homolog** | **spindle and kinetochore-associated protein 1 homolog** belongs to the spindle and kinetochore associated protein complex (Ska) is an essential component in chromosome segregation, but was only studied in mammals (Yu et al. 2021). |
| **Bhaga_2.g1623** | **protein IWS1 homolog** | Transcription factor which plays a key role in defining the composition of the RNA polymerase II (RNAPII) elongation complex and in modulating the production of mature mRNA transcripts (Liu et al. 2007). |
| **Bhaga_2.g1733** | **protein transport protein SEC16B homolog** | Protein transport protein Sec16B also known as is a protein that in humans is encoded by the SEC16B gene (NCBI 2025). |
| **Bhaga_2.g1756** | **CCR4-NOT transcription complex subunit 1** | Holoprosencephaly is the incomplete separation of the forebrain during embryogenesis, but was only studied in humans (Kruszka et al. 2019). |

**S15:** Coding region of the gene variant *Bhaga_2.g857* (*polygalacturonase QRT3-like*) and the significant marker on chromosome 2 at ~ 7.543 Mb (green).

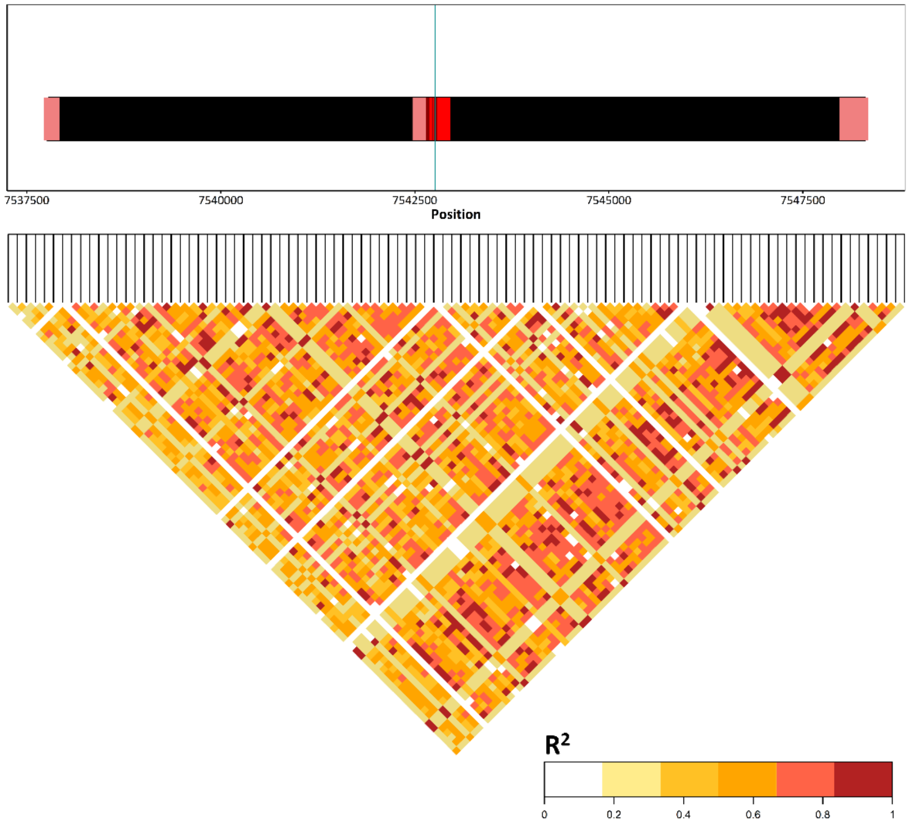


**S16:** Observed codon changes which lead to an amino acid (AA) replacement and a resulting change in polar bonds (hydrophobic or hydrophilic) at the underlying gene variants of markers associated with stomatal density

| **Gene variant** | **Variant version** | Position in bp | **Change in polar bonds** | **Original** | **Replaced by** |
|---|---|---|---|---|---|
| **Bhaga_2.g857** | missense variant | 7542764 | Hydrophilic AA was replaced by hydrophobic AA | Ala | Pro |
| **Bhaga_2.g883** | missense variant | 7787374 | Hydrophilic AA was replaced by hydrophobic AA | His | Asp |

**S17:** Coding region of the gene variant *Bhaga_2.g883* (*NRT1/PTR_FAMILY 5.4-like*) and the significant marker on chromosome 2 at ~ 7.787 Mb (green).

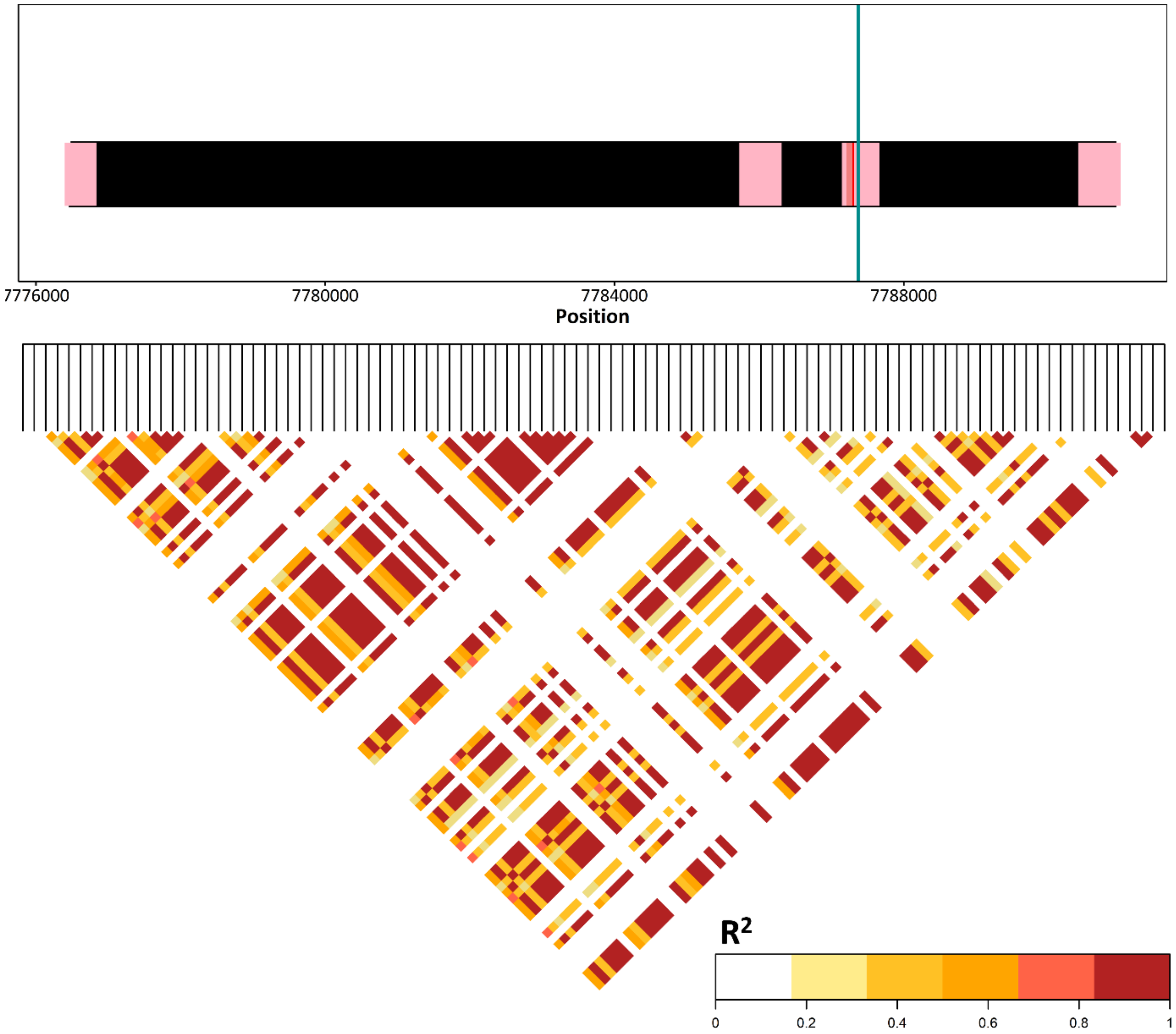
